# Contrasting Kinetics of Highly Similar Chloroalkane Reductive Dehalogenases

**DOI:** 10.1101/2024.07.10.602960

**Authors:** Katherine J. Picott, Elizabeth A. Edwards

## Abstract

Chloroform and trichloroethanes are pervasive groundwater contaminants for which bioremediation has been an effective treatment strategy. Reductive dehalogenase (RDase) enzymes from organohalide-respiring bacteria are essential for their remediation under anaerobic conditions. RDases are responsible for dehalogenating these chlorinated solvents, leading to their removal. This work explores the kinetic characteristics of three closely related *Dehalobacter* chloroalkane-reductases—TmrA, CfrA, and AcdA—and identifies differences between their activity on chloroform (CF), 1,1,1-trichloroethane (TCA), and 1,1,2-TCA. The side-by-side comparison of these enzymes has emphasized that TmrA and AcdA are specialized toward CF with both having 4-fold higher maximum specific activity (*V_max_*) on CF than 1,1,1-TCA, whereas, CfrA has very similar rates on both CF and 1,1,1-TCA. AcdA is the most sensitive to substrate inhibition by CF and 1,1,2-TCA, and inhibition by a common co-contaminant trichloroethene. Finally, the reduction of 1,1,2-TCA, which can produce both 1,2-dichloroethane and vinyl chloride, was assessed for each enzyme. Interestingly, each enzyme has a distinct preference for the major product it produces, indicating a favoured reaction pathway. Despite over 95% sequence identity, TmrA, CfrA, and AcdA exhibit substantial differences in kinetic behaviour, highlighting the importance of understanding such nuances for informed bioremediation strategies.

**SYNOPSIS:** Three similar dechlorinating enzymes were found to have distinct reaction rates and levels of sensitivity to inhibition. These differences have implications for the enzymes’ use in bioremediation.

## INTRODUCTION

Chloroform (CF) and isomers of trichloroethane (TCA) are widespread groundwater contaminants that are included on the Agency for Toxic Substance and Disease Registry’s (ATSDR) Substance Priority List ^1,2^. Bioaugmentation is a convenient and economical method of remediating sites contaminated with these compounds ^2–5^. There are several bioaugmentation cultures that target CF, 1,1,1-TCA, and 1,1,2-TCA ^6–11^.

These bioaugmentation cultures are all mixed communities that contain organohalide-respirers *Dehalobacter* and *Desulfitobacterium* ^7,8,11–14^, which use reductive dehalogenases (RDases) as the terminal enzyme in their electron transport chain ^15^. RDases catalyze the reduction of specific organohalide compounds—specificity depends on the enzyme itself—and in doing so, they remove one halogen through a hydrogenolysis pathway or two halogens through a dihaloelimination pathway ^15^. All RDases require a cobamide and two iron-sulfur clusters as cofactors ^15^. RDases with high protein sequence identity (>90%) are classified into ortholog groups (OGs), members of each OG generally target substrates that are structurally similar; e.g. chloroalkanes ^16,17^. However, it has been observed that similar RDases can be tuned to have uniquely specialized activity ^18–20^, yet the factors contributing to these differences are largely unexplored.

OG 97 comprises one *Desulfitobacterium* and six *Dehalobacter* RDases that dechlorinate halomethanes and chloroethanes ^7,9,10,20–22^. OG 97 members are particularly intriguing because they are the only identified RDases capable of dechlorinating CF and 1,1,1-TCA, and they display variation in substrate preferences despite many sharing over 95% amino acid sequence identity ^7,9,10,20–22^. The substrate preferences of the enzymes correlate well with the respiration of their native organism ^7,9,10,20–22^, indicating that the RDase activity influences respiratory function. Notably, *Dehalobacter* strains producing different RDases—TmrA, CfrA, AcdA, and RdhA D8M_v2_40029—exhibit varying CF isotope fractionation patterns, potentially indicating differences in the enzymatic mechanisms ^10,23,24^.

This study focuses on TmrA from *Dehalobacter* sp. UNSWDHB in CFH2 ^21,25^, CfrA from *Dehalobacter* sp. CF in ACT-3 ^20^, and AcdA from a *Dehalobacter* sp. in the enrichment SC05-UT—each found in deployed bioaugmentation cultures ^14,22^. RdhA D8M_v2_40029, which shows similarly suppressed carbon isotope fractionation patterns to TmrA and AcdA ^10,23,24^, was not included. Previous reports on TmrA, CfrA, and AcdA show similar activity on CF, 1,1,1-TCA, and 1,1,2-TCA; TmrA and AcdA also act on 1,1-dichloroethane (DCA), whereas CfrA does not ^20–22,26,27^. The kinetics of TmrA and CfrA have been evaluated using cell-free extracts and purified TmrA on select substrates ^6,21,28,29^. However, these studies were conducted under different conditions ^6,21,28,29^, limiting the ability to correlate kinetic data with the observed activity discrepancies. A side-by-side kinetic comparison with these purified chloroalkane reductases has not yet been conducted but could provide valuable insights into the catalytic differences among these closely related enzymes. Gaining a deeper understanding of RDase biochemistry benefits bioremediation practitioners by improving predictions of substrate degradation rates, byproduct formation, and potential inhibition.

The recent ability to heterologously express these enzymes in a common host enables a direct comparison ^26^. Here, we analyze the kinetics of heterologously produced TmrA, CfrA, and AcdA against three substrates: CF, 1,1,1-TCA, and 1,1,2-TCA. We also compare their sensitivity to inhibition by a common co-contaminant, trichloroethene (TCE), and examine structural features that could contribute to the observed differences.

## MATERIALS AND METHODS

### Materials

All chemicals, unless otherwise stated, BugBuster^®^, and filter tubes were purchased from MilliporeSigma (Burlington, MA, USA). All growth media, antibiotics, and induction agents were purchased from BioShop Canada Inc. (Burlington, ON, Canada). Bradford reagent was purchased from Bio-Rad Laboratories Ltd. (Hercules, CA, USA). All glass syringes were from Hamilton Company (Reno, NV, USA). Steel washers were purchased from Amazon.ca (ASIN: B081L91F6N). All gasses were supplied from Linde Canada Inc. (Mississauga, ON, Canada). A Coy vinyl anaerobic chamber was maintained with an anaerobic atmosphere of 10% CO_2_/2-3% H_2_/Bal. N_2_ (v/v) (Coy Laboratory Products Inc., Grass Lake, MI, USA); this is referred to as a glovebox throughout. Gas chromatography materials were purchased from Agilent Technologies (Santa Clara, CA, USA).

### Enzyme Expression

Three RDases were heterologously expressed in *Escherichia coli*: TmrA (accession: WP_034377773) from *Dehalobacter* sp. UNSWDHB ^21^, CfrA (AFV05253) from *Dehalobacter* sp. CF ^20^, and AcdA from a *Dehalobacter* in the SC05-UT enrichment culture (XCH81453) ^22^. The construction of the expression plasmids *p15TVL-tmrA*, *p15TVL-cfrA*, and *p15TVL-acdA,* enzyme expression, and partial purification have been described elsewhere ^22,26^. Minor modifications to the protocol are: lysogeny broth (LB) media was used, and Triton X-100 and glycerol were removed from the wash, elution, and storage buffers. ^22,26^. Each expression plasmid was expressed with *pBAD42-BtuCEDFB* in *E*. *coli* BL21(DE3) *cat-araC*-P_BAD_-*suf*, *ΔiscR::kan, ΔhimA::*Tet^R^ ^30,31^. The *pBAD42-BtuCEDFB* plasmid was gifted by the Booker Lab (Pennsylvania State University, PA, USA). The *E. coli* strain was gifted by the Antony and Kiley Labs (St. Louis University School of Medicine, MO, USA; University of Wisconsin-Madison, WI, USA). All plasmids are in Table S1 and a detailed methodology can be found in Supplemental Text S1.

The protein concentration was measured by Bradford assay ^32^, and the purity was estimated by sodium dodecylsulfate polyacrylamide gel electrophoresis (SDS-PAGE) using Image Lab 6.1 software (2022, Bio-Rad Laboratories, Inc.), (Figure S1). The standard deviation of triplicate protein concentration measurements was incorporated into the enzyme activity error (Table S2).

### Kinetic Experimental Method Design and Assumptions

The substrates and products in these reactions are volatile. During preliminary assay development (data not shown), we found that using bottles with headspace produced less consistent mass balances than glass gas-tight syringes without headspace. This is likely due to the short incubation time and frequent sampling invalidating the gas-liquid equilibrium assumption. The final iteration of the assay was conducted in the barrel of a 1-mL glass gas-tight Hamilton syringe. The total assay volume was sampled and analyzed by gas chromatography-flame ionization detection (GC-FID). Given the consistent mass balance we assume that no substrate or product is lost, and the initial amount of substrate was calculated from the amount of product and remaining substrate.

The elimination of the headspace, in turn, reduced mixing in the reaction, as observed with highly variable reaction rates in initial trials. We added a small stainless-steel washer inside the syringe barrel as a physical mixer (inspired by Criddle, 1989 ^33^) which improved consistency. We assume that these reactions are well-mixed and that the rates are governed by the enzymatic rate-limiting step.

Kinetic assays were conducted assuming the chlorinated solvent was the only limiting substrate. RDases depend on an external supply of electrons for catalysis. If the electron donor were limiting (i.e., not in excess of the target substrate), then the apparent rate would be underestimated.

Detailed descriptions of the assay setup, data analysis, and kinetic model assessment are provided in Supplemental Texts S2-4. It is important to note that in the native *Dehalobacter*, the rate-limiting step of respiration may be the supply of electrons through the electron transport chain rather than the RDase activity, as suggested previously ^29,34^. Our study focuses solely on the rate dictated by the dehalogenation of the supplied organohalide substrate. Additionally, the reactions were performed at room temperature, but the exact temperature could not be controlled in the glovebox and varied between 20–25°C (discussed further in Supplemental Text S5). We will only compare assays performed at the same recorded temperature.

### Kinetic Assay – Vial Set-up

All kinetic assays were performed in an anaerobic glovebox. All materials were equilibrated in the glovebox atmosphere for at least 48 hr before use. A series of vials were used to prepare the reaction buffer for 15 different substrate concentrations with one substrate-free vial to produce a full Michaelis-Menten curve. The reactions themselves took place in glass syringes— the prepared buffer was taken from the vial and the desired enzyme was added to the syringe afterward to keep the stock buffer clean. Four reactions total were done using the same stock buffer vial (enzyme-free negative control, TmrA, CfrA, and AcdA).

The assay buffer and substrate mixtures were prepared in sixteen 11-mL vials with mininert caps. Each vial was prepared so that the final liquid volume would be 6 mL with 5 mL headspace. Reaction buffer containing 50 mM Tris-HCl (pH 7.5) and 1 mM methyl viologen (MV) was added to each of the vials, the vials were capped, and the substrate was delivered to the desired concentration from either a saturated water stock or the neat substrate using a gas-tight glass syringe (Table S3). Target aqueous substrate concentration ranges were 10 µM–1.5 mM chloroform (CF), 5 µM–2 mM 1,1,1-trichloroethane (TCA), and 10 µM–5 mM 1,1,2-TCA. However, the actual initial substrate concentration was calculated by summing the remaining substrate and the product measured by GC-FID. All vials were equilibrated overnight. After equilibration, the reductant Ti(III) citrate (pH 7) was added to each vial to a final concentration of 5 mM. This was added directly before the assays to prevent premature oxidation of the reductant. Further details of the reaction vial setup and the assay procedure can be found in Supplemental Text S2.

Trichloroethene (TCE) was tested as an inhibitor for CF dechlorination. Glass bottles (255 mL) with mininert caps were set up with 60 mL of the buffer (50 mM Tris-HCl [pH 7.5] and 1 mM MV). TCE was delivered to a concentration of 50 µM using a TCE-saturated water stock. The stock solution was equilibrated overnight and then aliquoted to the 11-mL reaction mixture vials to all have the same amount of TCE. The substrate, CF, and Ti(III) citrate were delivered into the vials as previously stated.

### Kinetic Assay – Reaction

Purified RDases were thawed on ice and diluted in 50 mM Tris-HCl (pH 7.5) to an RDase concentration of 100 nM. Four—one for each enzyme and an enzyme-free control—1-mL glass, gas-tight Hamilton syringes with a flat M1.6 (4 mm diameter, 0.3 mm thick) stainless steel washer inside the barrel of the syringe were used to perform the reactions. The reactions were performed by taking up 900 µL of the assay buffer from the prepared reaction mixture vial and then 100 µL of the diluted enzyme, or buffer for the enzyme-free controls. The syringe was inverted continuously during the incubation, allowing the washer to move through the barrel and physically mix the solution. The reactions were run for 2 min for CF and 1,1,1-TCA assays, and 5 min for the 1,1,2-TCA assays. At the end of the incubation, the 1-mL reaction was injected via a submerged needle into 5 mL HCl acidified water (pH <2) in an 11-mL GC headspace autosampler vial and immediately sealed by crimping with a Teflon-coated septum. The assays were always carried out in order of the lowest substrate concentration to the highest concentration. The same syringe was used for each enzyme or control and washed between reactions with 1 mL of anaerobic acidified water and 1 mL of anaerobic double-deionized water. A schematic of the reaction procedure is in Figure S2.

The 1,1,1-TCA kinetic curve was constructed using two sets of data and outliers were identified and removed. Outliers occurred when 1,1,1-TCA was delivered using larger volumes (>250 µL) of saturated stock which caused depletion of the electron donor. The outliers were identified visually by the oxidation of methyl viologen in the buffer vial and if 1,1-dichloroethene (DCE) was present in the negative controls. 1,1,1-TCA is unstable in water and can undergo β-elimination to form 1,1-DCE and hydrolysis to form acetic acid ^35,36^, which does not interfere with the assay, but 1,1-DCE presence correlated with the observed abiotic oxidation. In these cases, the catalysis is limited by electron supply and the enzyme rate is underestimated. Increasing the number of reactions performed for 1,1,1-TCA and using fresh substrate stocks gave more confidence in the model fit despite the uneven coverage along the Michaelis-Menten curve.

All raw data (including outliers) for each kinetic trial is available in Table S4. Negative control data and raw product masses for each reaction are in Supplemental Text S6.

### Analytical procedures (gas chromatography)

Analysis of substrates and products was carried out by GC-FID following the same methods as previously described ^22,26^, full details are in Supplemental Text S2.

### Data processing

The sum of the remaining substrate and products was used to determine the initial substrate concentration in each reaction. The rate was determined by dividing the amount of product (nmol or mg) by the assay time (s) and the mass of RDase in the assay (mg). Error in the protein concentration and an estimated 2-3 s variance for the incubation time were used as uncertainty in the rate values (Table S4).

### Kinetic model fitting

The kinetic data were plotted in Python3 and fit with various Michaelis-Menten kinetic models with and without inhibition using the LMFIT package ^37^. The equations of the models tested and their assessment are described in Supplemental Texts S3 and S4. Models were produced using the least squares method and included experimental error in the residual calculation. Data for CF, 1,1,1-TCA, and 1,1,2-TCA without external inhibitors were fit to a typical Michaelis-Menten kinetic expression with and without a term for substrate inhibition. The presence of substrate inhibition was determined by the model of best fit. The best-fit model was selected based on reducing the R^2^, Akaike Information Criterion, and the standard deviation of the residuals. The kinetic parameters *V_max_* (nmol/s/mg), *K_M_* (µM), and *K_I,sub_* (µM; if applicable) were extracted from the models. The *k_cat_* (s^−1^) was calculated by multiplying the *V_max_* by the mass of the protein (48 kDa; 48 000 g/mol) to convert the protein mass from mg to nmol. All calculated kinetic parameters— in µM and mg/L units—and fit evaluations can be found in Tables S5-S7.

To determine the inhibition constant, *K_I,inh_,* of TCE, first, the *V_max_*, *K_M_*, and *K_I,sub_* (if applicable) values were determined by the uninhibited CF assay performed on the same day. These values were then set as constants and the TCE-inhibited kinetic data was fit with models for competitive, non-competitive, and uncompetitive inhibition. The best-fit model—selected using the same criteria above—was used to estimate the *K_I,inh_*. All model assessments can be found in Table S8.

Kinetic analysis of 1,1,2-TCA was performed differently than the other substrates because two products are formed (scheme in Figure S3B). The *K_M_* and *K_I,sub_* were calculated by modelling the overall 1,1,2-TCA consumption. The reaction rates for each product were then determined by setting *K_M_* and *K_I,sub_* as the values determined from the overall rate. A detailed discussion about the 1,1,2-TCA kinetic analysis can be found in Supplemental Text S3.

### Protein Models

Protein models of TmrA and CfrA were retrieved from AlphaFill ^38^. AlphaFill contains a database of AlphaFold2 models of all sequences available in UniProt and docks cofactors into the models based on homology to known crystal structures ^38,39^. Structures for TmrA and CfrA with cobalamin and two iron-sulfur clusters were extracted and further relaxed using YASARA energy minimization server ^40^. The AcdA model was generated directly from AlphaFold2 without the cofactors. The crystal structure of PceA (PBD ID: 4ur0) was used as a comparison of the conserved active site residues ^41^. All structures were visualized and analyzed using PyMOL Molecular Graphics System v2.3.4 ^42^.

## RESULTS & DISCUSSION

### Activity on Chloroform and 1,1,1-Trichloroethane (Methyl Chloroform)

Despite CF and 1,1,1-TCA differing by only one methyl group, RDase activity varies significantly between these compounds. Figure 1 and Table 1 show that TmrA and AcdA have much higher *V_max_* and lower *K_M_* values with CF than 1,1,1-TCA. Conversely, CfrA dechlorinates CF and 1,1,1-TCA as substrates at similar rates, but still with a lower *K_M_* for CF. This affinity trend aligns with previously published data despite the greater absolute values (Table 1) ^29,43,44^. The maximum rates observed for the purified enzymes are 500–100-fold greater than the literature values. Differences in assay setup and the catalyst could cause this. Most of the previous data was performed with cell-free lysates ^6,28,29,44^, in which the RDase concentration is unknown, causing underestimations of the rate. Additionally, there may be protein-protein interactions in the cell lysate that alter protein-substrate affinity not captured in pure protein. TmrA purified from *Dehalobacter* also had a lower affinity and rate ^21^. The reaction time may also cause these discrepancies—previous data was measured from 1-5 h incubations ^6,21,28,44^. A long incubation time underestimates the rate because the substrate and electron donor are depleted over time causing the RDase rate to decrease from its initial rate.

**Figure 1.**
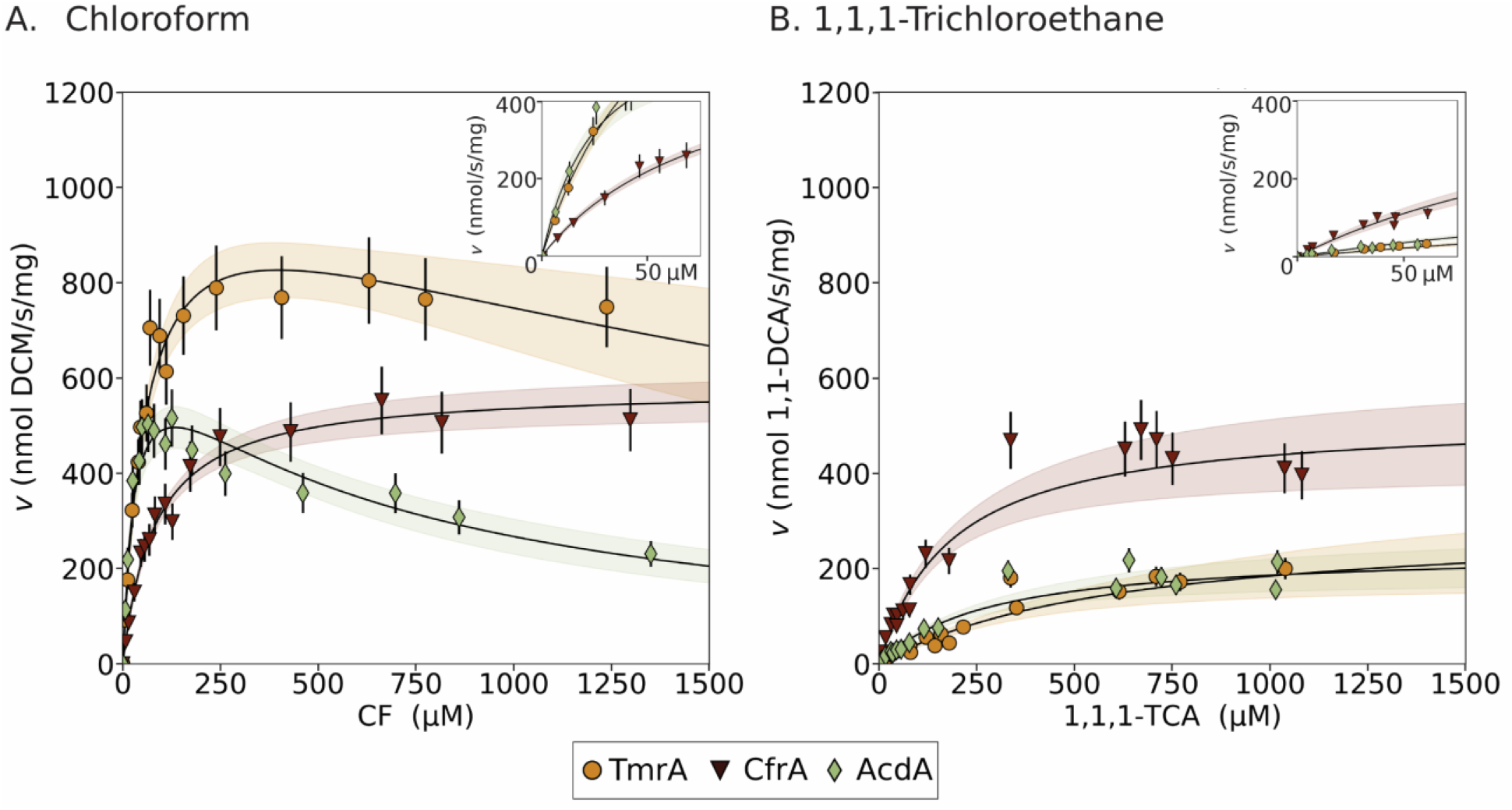
Kinetic curves for TmrA (yellow circles), CfrA (red triangles), and AcdA (green diamonds) using (A) chloroform (CF) and (B) 1,1,1-trichloroethane (1,1,1-TCA) as substrates. Inset plots show the rate of reaction for substrate concentrations lower than 70 µM. Error bars represent the uncertainty in the calculated rate. The black lines are the optimized kinetic models, shaded area represents the 95% confidence interval. All reactions were 2 min at 24°C.

**Table 1.**
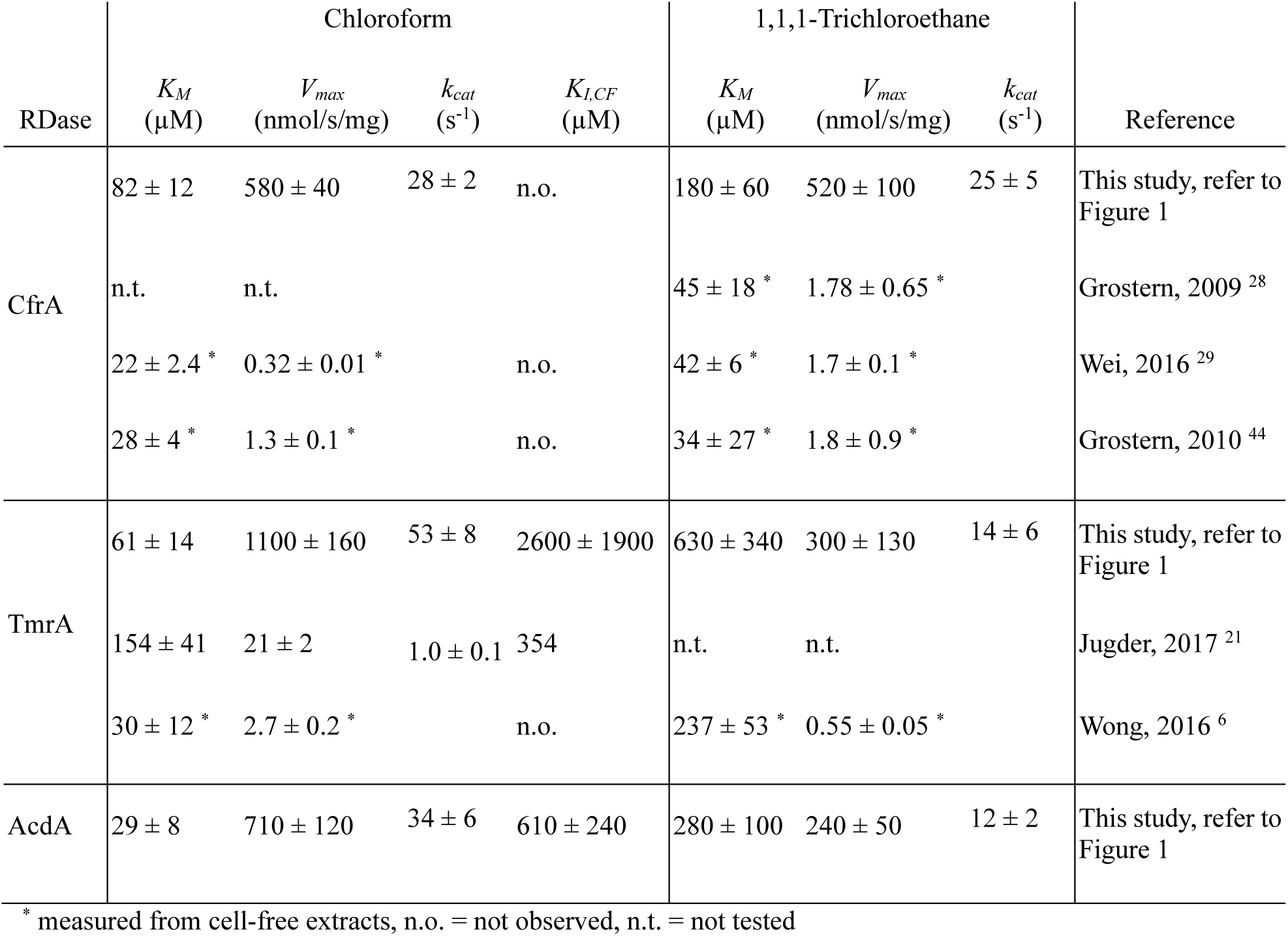
Kinetic parameters with chloroform and 1,1,1-trichloroethane, error is the 95% confidence interval.

TmrA and AcdA’s higher affinities for CF than 1,1,1-TCA suggest that they have a more compact active site than CfrA, whose difference in *K_M_* is not as drastic. Although, at high concentrations of CF, TmrA and AcdA experience substrate inhibition. Substrate inhibition occurs when the substrate binds unproductively and prevents catalysis, thereby reducing enzyme activity as substrate concentration increases. A low *K_I,sub_* corresponds to stronger inhibition because the value represents the concentration needed to reduce the enzyme to half of its maximum rate. This principle is the same for external inhibitors. With a lower *K_I,CF_*, AcdA experiences stronger substrate inhibition, but at concentrations of CF sub 100 µM the rate is comparable to TmrA (Figure 1A).

In published data, cell-free extracts containing CfrA were observed to have inhibition by 1,1,1-TCA with a *K_I,sub_* range of 300–1000 µM ^28,44^. In this work, efforts to fit the 1,1,1-TCA kinetic data with a substrate-inhibition model were unsuccessful, likely due to variance in reaction rate. This variance could be caused by electron donor consumption by 1,1,1-TCA abiotic degradation products, or by the low solubility of 1,1,1-TCA ^35,36^. 1,1,1-TCA has a low solubility (∼9 mM) in water ^45^, and though the tested concentrations are within this threshold, the buffer could interfere with the solubility reducing the concentration available for reaction. The cell-free extract experiments could have suffered from the same confounding variables.

### Trends of Activity on Native Substrates and Correlation with Kinetic Isotope Effects

The kinetic trends observed for each RDase correlate to the substrate structural differences and the enrichments from which the RDase was identified. The RDases generally exhibit higher *V_max_* and lower *K_M_* values with CF, likely due to CF’s smaller size facilitating better fit into the active site and faster reaction rates. Both TmrA and AcdA were identified from *Dehalobacter* originating from CF-contaminated sites ^12–14,22^, while CfrA comes from a *Dehalobacter* enriched from a 1,1,1-TCA and TCE-contaminated site ^11^. CfrA, having been the only of these exposed to 1,1,1-TCA, may have evolved a specialization resulting in a higher rate than the other two.

We can compare these kinetic trends with isotope effects, represented by the enrichment factor (ε). ε reflects how substrates with light isotopes (C^12^/Cl^35^) in reactive bonds (those forming or breaking) react faster, enriching the substrate pool with heavier isotopes (C^13^/Cl^37^) ^46^. These three enzymes are known to produce distinct isotope fractionation patterns when tested *in vivo* ^23,24^. Table 2 shows published values of ε for each enzyme—negative values indicate the enrichment of the heavy isotope; a positive value is an inverse isotope effect when the light isotope is enriched ^23^. Differences in isotope fractionation can occur due to mechanistic differences in the catalytic slow-step or from “masking effects”, which are slow pre-transformation steps that cause suppression of the isotope fractionation. Commitment to catalysis—where substrate binding and catalysis are much faster than substrate dissociation, reducing isotope discrimination—is one such effect (explained further here ^23,46,47^).

**Table 2.** The most recently published carbon and chlorine fractionation values for chloroform and the catalytic efficiency measured in this study; error is the 95% confidence interval.

| Catalyst<br>(culture) | $\epsilon^{13}\text{C}$ | $\epsilon^{37}\text{Cl}$ | $k_{cat}/K_M$<br>($\text{s}^{-1} \mu\text{M}^{-1}$) | Reference for<br>epsilons |
| --- | --- | --- | --- | --- |
| Abiotic |  |  |  |  |
| CNCbl | $-26.04 \pm 0.91\text{‰}$ | $-4.00 \pm 0.2\text{‰}$ | | Heckel, 2019<sup>23</sup> |
| CfrA |  |  |  |  |
| (ACT-3) | $-26.8 \pm 3.2\text{‰}$ | $-6.86 \pm 0.35\text{‰}$ | $0.34 \pm 0.06$ | Phillips, 2022<sup>24</sup> |
| TmrA |  |  |  |  |
| (CFH2) | $-3.1 \pm 0.5\text{‰}$ | $2.5 \pm 0.3\text{‰}$ | $0.87 \pm 0.24$ | Heckel, 2019<sup>23</sup> |
| AcdA |  |  |  |  |
| (SC05) | $-1.52 \pm 0.34\text{‰}$ | $-1.84 \pm 0.19\text{‰}$ | $1.2 \pm 0.4$ | Phillips, 2022<sup>24</sup> |

CNCbl = cyanocobalamin

The variance in ε values observed for CfrA, TmrA, and AcdA may result from different transformation mechanisms or structural changes causing masking ^24^. Although we cannot determine mechanisms from this kinetic analysis, we can compare catalytic efficiency (*k_cat_/K_M_*). The *k_cat_/K_M_* reflects the enzyme’s maximum rate and binding affinity. Generally, the *k_cat_/K_M_* increases in the same order that the ε^13^C decreases (Table 2). CfrA has similar fractionation values to the abiotic reaction catalyzed by reduced cobalamin, but TmrA and AcdA have high suppression ^23,24^. This suppression may result from higher commitments to catalysis, given that they have higher *k_cat_/K_M_* values than CfrA. This trend is less clear in the ε^37^Cl values due to TmrA’s inverse fractionation. Structural diversity between the RDases has been thought to cause these fractionation differences and would also impact their catalytic efficiency ^24^.

### Trichloroethene Inhibition

The inhibition of chloroethenes on chloroalkane reductive dechlorination and vice versa is well established at both the organism and enzyme level ^11,28,48–50^. For example, CF will potently inhibit each dechlorination step of TCE by *Dehalococcoides* ^29,49,50^; meanwhile, TCE and its dechlorinated products inhibit CfrA activity on CF ^28,29^. This reciprocal inhibition may cause a stalemate where no dechlorination occurs. To explore this interaction on other chloroform RDases, we assessed the inhibition of CF-dechlorination in the presence of ∼45 µM TCE, simulating conditions where TCE dechlorination is stalled. Although this is not an exhaustive assessment of inhibition, thus the *K_I,TCE_* values are less certain, it provides perspective into the differences in sensitivity between the three RDases to external inhibition.

All three enzymes showed better model fits with uncompetitive inhibition, as was previously reported for CfrA ^29^. AcdA had the most drastic decrease in apparent activity (Figure 2); CfrA and TmrA had similar inhibition. CfrA’s *K_I,TCE_* value was higher but in the same magnitude as that measured in cell-free extracts of *K_I,TCE_* of 40 ± 3 µM indicating that this assessment provides a good estimate of inhibition ^29^. While TCE inhibition is expected, the inhibition of AcdA at lower TCE concentrations than TmrA and CfrA should be considered when deployed in the field. Additionally, full inhibition studies should be conducted to gain more confidence in the *K_I,TCE_* values and assess inhibition by other co-contaminants.

**Figure 2.**
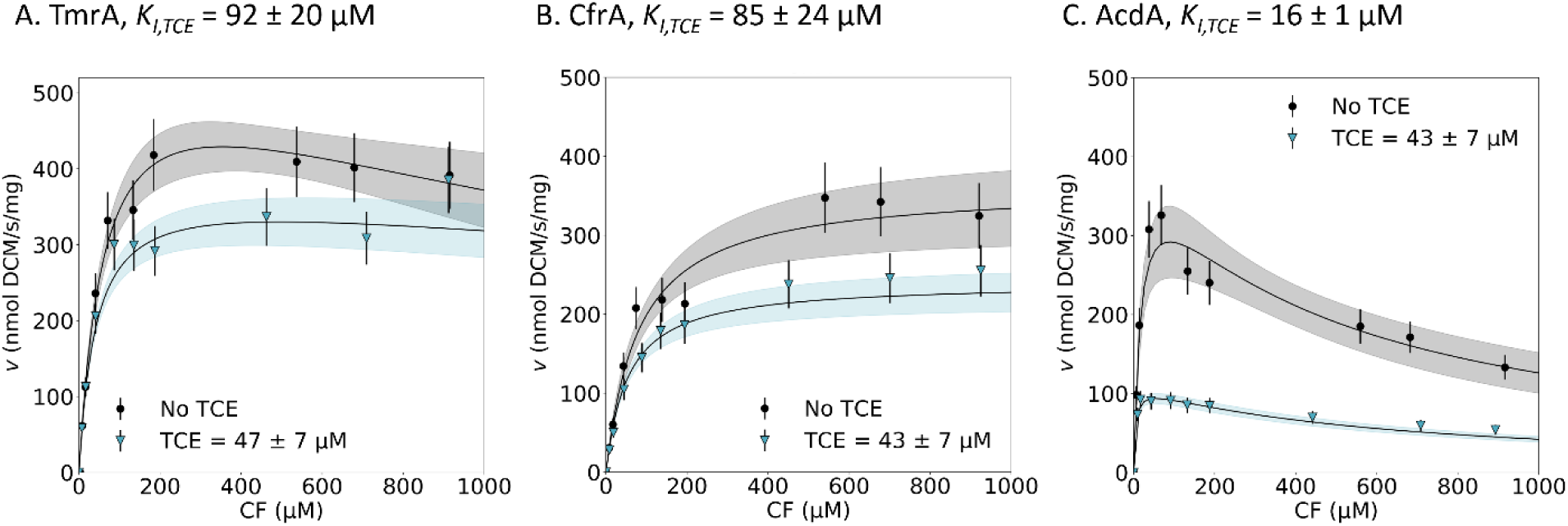
Kinetic curves for TmrA (A), CfrA (B), and AcdA (B) using chloroform (CF) as a substrate with trichloroethane (TCE) inhibition (blue triangle) and without (black circle). The average concentration of TCE between all reactions is presented on each plot, the error represents the standard deviation. Error bars represent the uncertainty in calculated rate. The black lines show the optimized kinetic model, shaded area represents the 95% confidence interval. All reactions were 2 min at 20°C.

### Kinetics on 1,1,2-Trichloroethane

1,1,2-TCA is a peculiar substrate because it can undergo both the typical hydrogenolysis to form 1,2-DCA or dihaloelimination to form vinyl chloride (VC). The dihaloelimination of 1,1,2-TCA is strongly favoured as it has a much higher redox potential (E_0_’ = 1063 mV, H_2_ as donor) than the hydrogenolysis reaction (862 mV) ^51^. Dihaloelimination can occur abiotically by reduced cobalamin at room temperature and neutral pH ^26^, it has also been observed by many RDases of varying specificities and likely occurs easily once the substrate enters the active site ^19,26^. Enzymes in OG 97 are known to catalyze both reactions, but the product of 1,1,2-TCA dechlorination can be consequential as VC is of higher regulatory concern and inhibits the dechlorination of co-contaminants CF and 1,1,1-TCA ^28,29^.

TmrA has the fastest turnover rate of 1,1,2-TCA, with over double the *V_max_* of CfrA and AcdA (Figure 3A, Table 3). AcdA shows the strongest substrate affinity, while CfrA has the largest *K_M_*, nearly ten times higher than AcdA. Regardless of the overall transformation of 1,1,2-TCA, it is important to look at the production rate of the two products, 1,2-DCA and VC, separately. TmrA still achieves the fastest rate of hydrogenolysis to 1,2-DCA (Figure 3B, Table 3). Both TmrA and AcdA perform dihaloelimination to produce VC with similar *V_max_* values, but AcdA slightly favours this reaction over hydrogenolysis, resulting in VC as the major product. In contrast, CfrA produces negligible amounts of VC. This homogenous production of 1,2-DCA simplifies subsequent remediation steps and minimizes VC inhibition of other dechlorination processes. Despite its slower rate under field-relevant conditions, CfrA exhibits the most desirable activity against 1,1,2-TCA.

**Figure 3.**
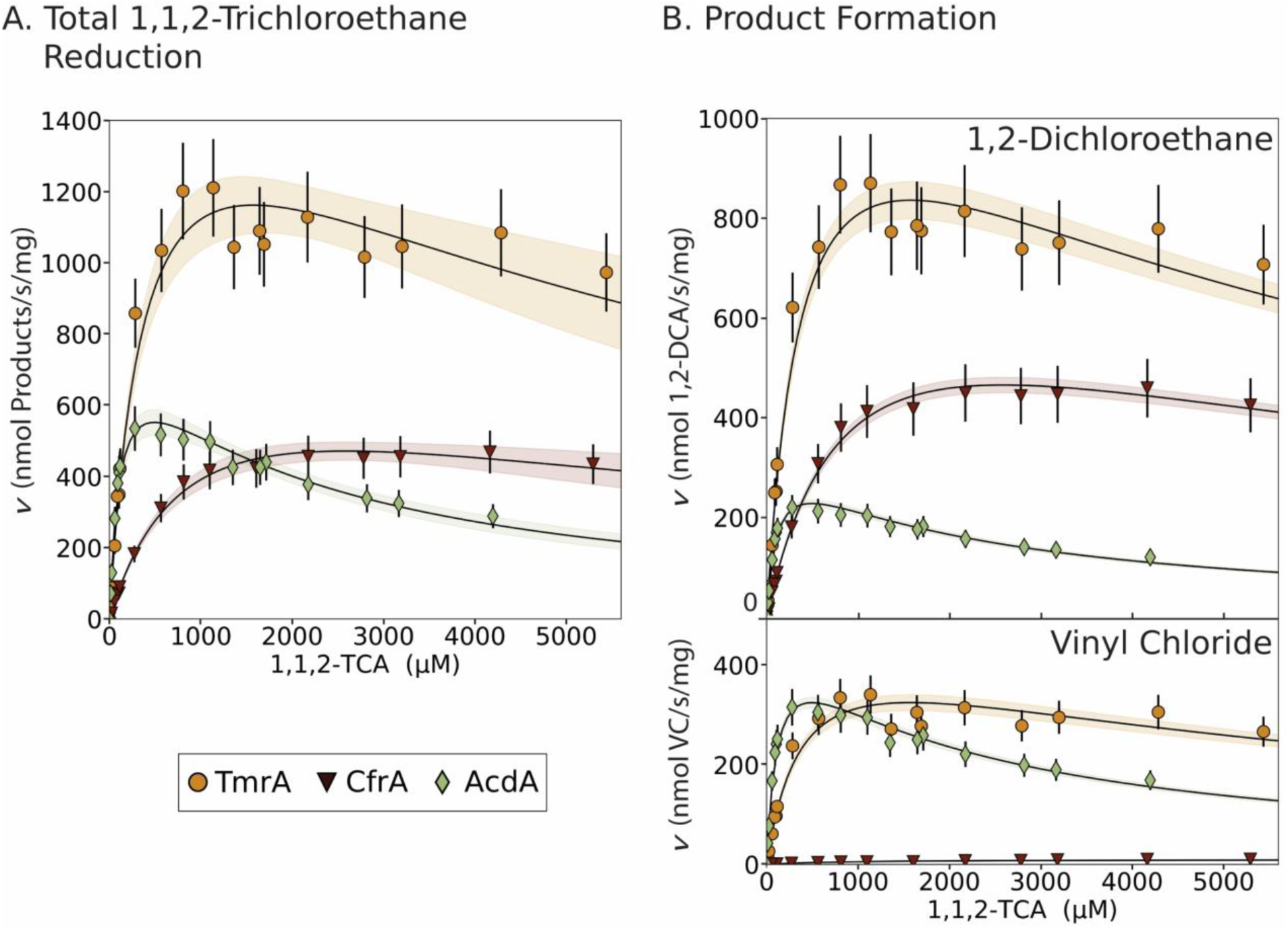
Kinetic curves for TmrA (yellow circles), CfrA (red triangles), and AcdA (green diamonds) using 1,1,2-trichloroethane (1,1,2-TCA) as a substrate. (A) is the total reduction of 1,1,2-trichloroethane and (B) is the production of 1,2-dichloroethane (1,2-DCA; top) and vinyl chloride (VC; bottom). Error bars represent the propagated uncertainty in the reaction rate. The black lines show the optimized kinetic model, shaded area represents the 95% confidence interval. All reactions were 5 min at 24°C.

**Table 3.** Kinetic parameters for 1,1,2-trichloroethane as a substrate resulting in either 1,2-dichloroethane or vinyl chloride as products, error is the 95% confidence interval.

| RDase | $K_M$<br>($\mu\text{M}$) | $K_{I,TCA}$<br>($\mu\text{M}$) | Total<br>$V_{max}$<br>(nmol/s/mg) | Total<br>$k_{cat}$<br>($\text{s}^{-1}$) | 1,2-DCA<br>$V_{max}$<br>(nmol/s/mg) | 1,2-DCA<br>$k_{cat}$<br>($\text{s}^{-1}$) | VC<br>$V_{max}$<br>(nmol/s/mg) | VC<br>$k_{cat}$<br>($\text{s}^{-1}$) | Percent of<br>1,2-DCA<br>in product<br>(%) |
| --- | --- | --- | --- | --- | --- | --- | --- | --- | --- |
| CfrA | 950 ± 220 | 7100 ± 4400 | 820 ± 160 | 39 ± 4 | 810 ± 20 | 38 ± 0.5 | 10 ± 0.6 | 0.48 ± 0.01 | 98.7 ± 0.6 |
| TmrA | 400 ± 96 | 6200 ± 3700 | 1800 ± 320 | 86 ± 8 | 1300 ± 50 | 62 ± 1 | 490 ± 20 | 24 ± 0.5 | 72.0 ± 1.1 |
| AcdA | 120 ± 20 | 2000 ± 550 | 820 ± 100 | 39 ± 2 | 340 ± 10 | 16 ± 0.5 | 480 ± 15 | 23 ± 0.5 | 41.2 ± 0.5 |

TCA = trichloroethane, DCA = dichloroethane, VC = vinyl chloride

Interestingly, the product composition remains consistent for each enzyme across 1,1,2-TCA concentrations and incubation times (Figure S11, Table 3). CfrA produces 98.7 ± 0.6% 1,2-DCA; consistent with 98.8 ± 0.5% previously measured from cell-free extracts ^26^. The 1,2-DCA composition produced by TmrA (72 ± 1%) and AcdA (41.2 ± 0.5%) is within 4% of published values performed with longer incubation times ^22,26^. This consistency across varying conditions suggests the product distribution is inherent to each OG 97 enzyme and potentially predictable based on structural features controlling it.

### Variance In Catalytic Residues

Despite over 95% amino acid sequence identity equating only 15–21 amino acid differences (Supplemental Text S7), CfrA, TmrA, and AcdA exhibit distinct kinetic characteristics. Since these enzymes were all expressed heterologously, these differences must be attributed to the peptide sequence rather than variations within the actual organism—such as the type of cobamide used, which can affect the dechlorination activity ^52–54^. Although experimentally derived structures of chloroalkane reductases remain elusive, protein modelling offers some insights. The resolution of the PceA (*Sulfurospirillum multivorans*) crystal structure revealed several highly conserved active site residues critical for activity ^41,55^. Many differences we observe between the OG 97 RDases overlap or collocate with these important residues. The identity of residues involved in protonation during hydrogenolysis is of particular interest as they directly impact the dehalogenation reaction and, therefore, the rate.

PceA features a tyrosine (Tyr246) at the substrate binding site, long thought to be the primary proton donor due to its position and necessity for enzyme activity and conservation in many RDases including those from *Dehalococcoides* and *Dehalobacter* (Supplementary Text S7) ^19,41^. Mutagenic studies also implicated a neighbouring asparagine Asn272 and arginine Arg305 in catalysis ^19,41,55^, believed to stabilize Tyr246 via H-bonding, evident from their proximity in crystal structures (∼3 Å; Figure 4A). However, a recent computational study by Zhang *et al.* suggests the more energetically favourable proton donor is Arg305, with Tyr246 and Asn272 positioning Arg305 toward the substrate ^56^.

**Figure 4.**
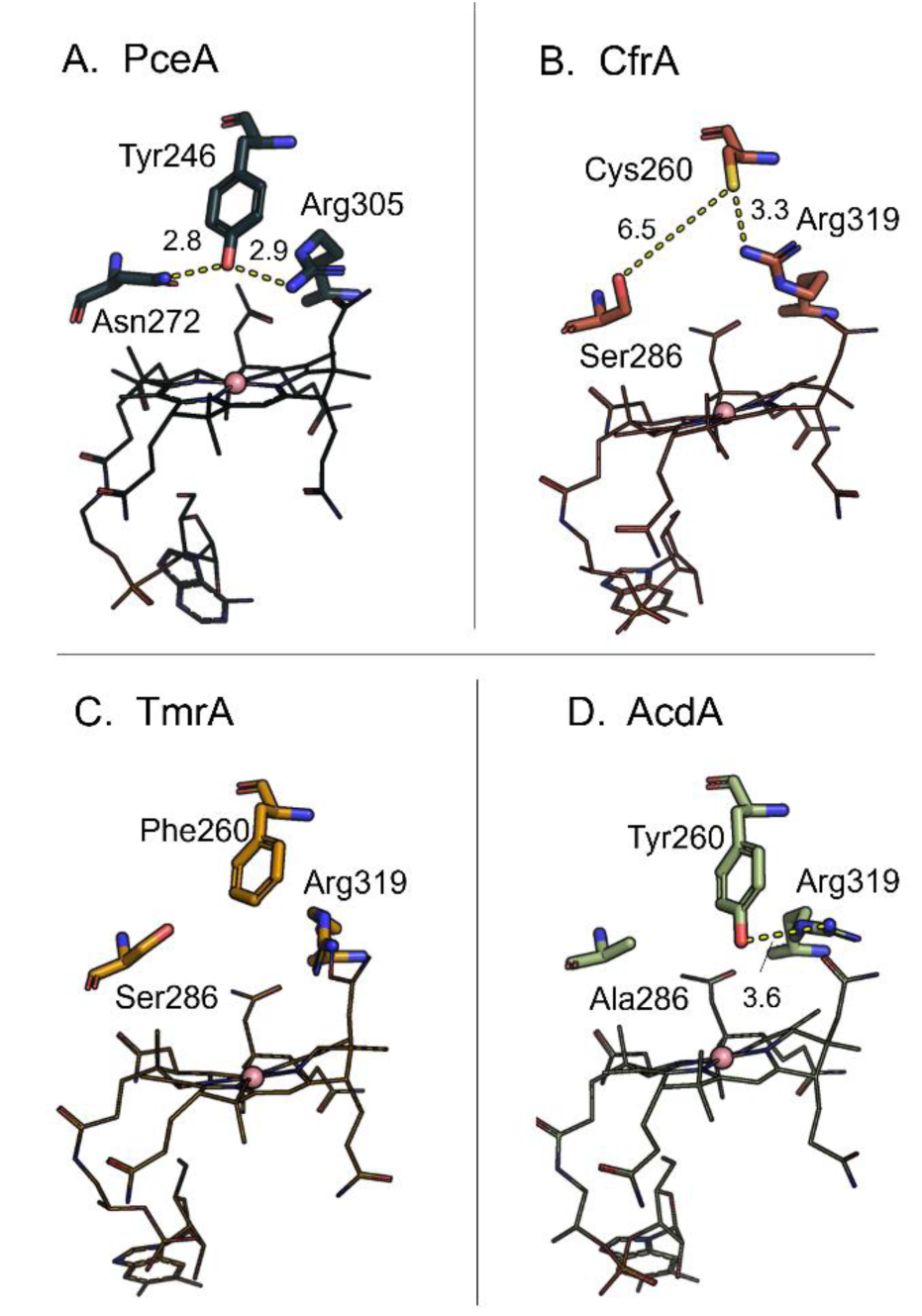
The conserved active site residues in the PceA crystal structure with predicted catalytic activity and the corresponding residues in CfrA, TmrA, and AcdA protein models. Distances between key residues are shown as dashed lines with the distance in angstroms. Wire structure is the cobamide cofactor and the pink sphere is the cobalt. PDB ID: 4ur0 ^41^.

This hypothesis better explains the activity of CfrA and TmrA, which lack tyrosine at the expected position but have the arginine Arg319 (Figure 4B/C). TmrA instead has a non-H-bonding phenylalanine (Phe260) and CfrA has a cysteine (Cys260), which may have a weaker H-bond due to the distance (3.3 Å in the model) and leaves a larger space in the active site. Both enzymes have serine (Ser286) instead of Asn272, capable of H-bonding. AcdA retains the conserved tyrosine (Tyr260) and arginine (Arg319) but has alanine (Ala286) instead of asparagine (Figure 4D). This asparagine is less conserved in RDases (Supplemental Text S7), so its exact role is unclear, but a network of interactions could potentially affect the catalytic residue positioning relative to substrates.

Regardless of their high sequence identity, TmrA, CfrA, and AcdA have drastic differences in these ostensibly highly conserved residues that likely influence their kinetic properties. For instance, an aromatic residue in position 260 may support a tighter interaction with the substrates leading to TmrA and AcdA having lower *K_M_* values for CF and 1,1,2-TCA than CfrA (more detail in Supplemental Text S7). Phillips *et al.* also identified other residues (Tyr80, Leu388, and Met391 in CfrA; Phe80, Phe388, and Trp391 in TmrA and AcdA) as possible sources of isotopic fractionation variance ^24^. Future mutant studies should explore these catalysis-implicated residues to understand their impact on kinetic characteristics and isotope effects.

### Implications For Bioremediation

This is the first true comparative kinetic study of highly similar purified RDases, revealing that even near-identical RDase variants can exhibit distinct behaviours that could influence remediation processes. First, kinetic parameters can be incorporated into groundwater models to help interpret field data and improve predictions of remedial progress. For example, establishing enzymatic rates can be useful for monitoring natural attenuation. Recent studies used quantitative PCR and proteomics to quantify active RDases in aquifers ^57,58^. By combining RDase and substrate concentrations with kinetic parameters, degradation rates (yr⁻¹) can be predicted using the Michaelis-Menten model ^57,58^. Predicting rates is necessary for decision-making at contaminant sites, and expanding the library of RDase kinetic coefficients will facilitate the implementation of this technique across more compounds.

Understanding the link between RDase sequence and associated differences in activity in terms of substrate preferences, products and inhibitors will aid in the interpretation field data at contaminated sites. Sequence variants of closely related RDases are sometimes detected at field sites, but their impact is generally not considered in interpreting results; the published features of the RDase is assumed without consideration to possible differences due to sequence variants. However, as molecular tools become more accessible, sequence variants will be more easily detected. For example, sequence variants within OG 97 were detected at a 1,1,1-TCA-contaminated site and shown to correlate with activity on either 1,1,1-TCA or 1,1-DCA substrates^59^. The different rates and sensitivity to common inhibitors found here likely extend to other OGs, highlighting the need for further comparative studies. Knowledge of RDase variant kinetics and inhibition could explain why natural attenuation sometimes stalls despite the presence of dechlorinators and appropriate RDases. If inhibition is an issue, biostimulation strategies would be ineffective, requiring alternatives for alleviating the inhibition ^50^.

Furthermore, this work outlines an accurate syringe barrel method for directly assay enzyme kinetics, expediting inhibitor screening, and revealing functional differences between RDases and organohalide respirers. Investigating the structural basis for kinetic variation will elucidate significant active site residues and guide future selection or engineering of more efficient RDases.

## ACKNOWLEDGEMENTS

This study was supported by the Government of Ontario with an Ontario Graduate Scholarship to K.J.P, by the Natural Science and Engineering Research Council (NSERC) with a Discovery Grant to E.A.E and a Canadian Research Chair to E.A.E. We thank the Booker Lab (Pennsylvania State University), the Antony Lab (St. Louis University School of Medicine), and the Kiley lab (University of Wisconsin-Madison) for gifting materials.

## ASSOCIATED CONTENT

### Supporting Information

Detailed methods for RDase production, additional assay details, and equations and explanations for all kinetic models used, discussion about the temperature dependence observed, graphical representations of the raw data and substrate conversion, and amino acid sequence alignments of RDases to complement structural comparison. (PDF)

All raw data for the assays run, including the negative controls, and all of the parameters and statistical values for each model fitting. (XLSX)

## AUTHOR INFORMATION

### Author Contributions

E.A.E. and K.J.P. conceived of the project. All experiments and data analysis were carried out by K.J.P. The manuscript was written through contributions of all authors. All authors have given approval to the final version of the manuscript.

### Notes

The authors declare no competing financial interest.

## TOC FIGURE

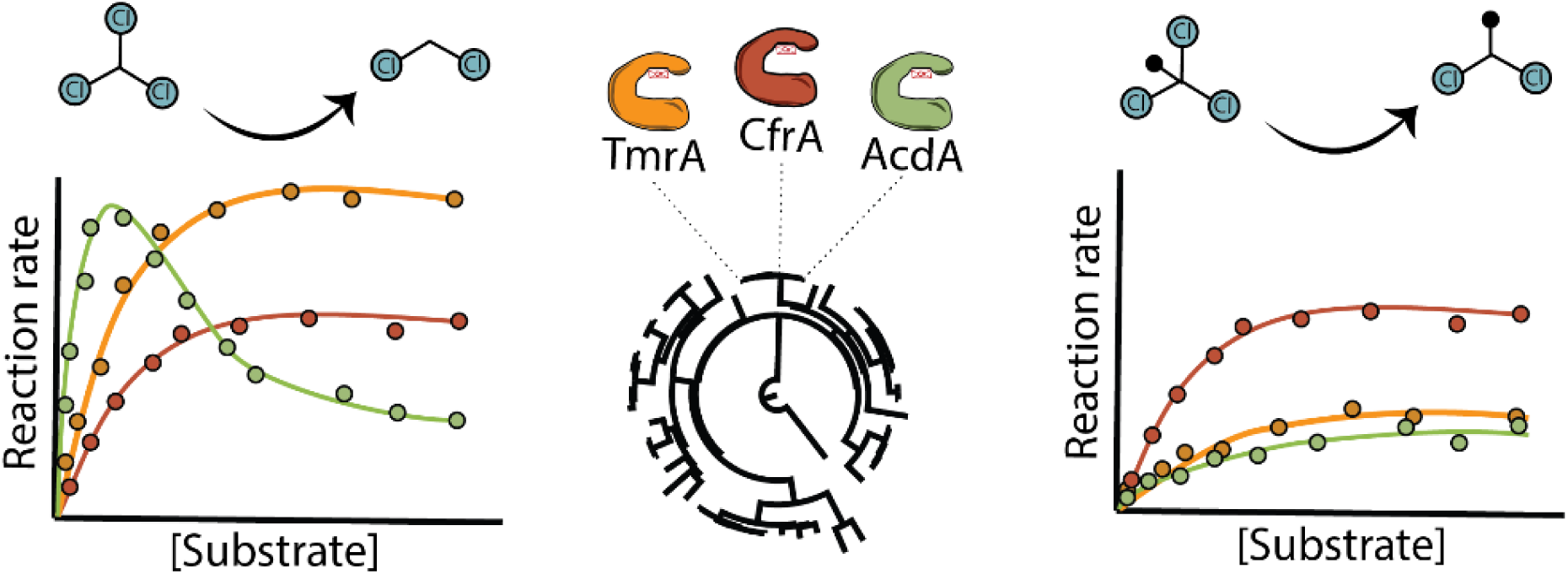

