## Supplemental Information for "Contrasting Kinetics of Highly Similar Chloroalkane Reductive Dehalogenases"

**Running Head:** Chloroalkane RDase Kinetic Comparison

#### SUPPLEMENTAL TEXT

| | |
|---|---|
| **Text S1 –** RDase Expression and Purification | …………………………S3 |
| **Text S2 –** Detailed Kinetic Assay Methods | …………………………S5 |
| **Text S3 –** Enzyme Kinetic Models | …………………………S9 |
| **Text S4 –** Raw Data and Model Fits | …………………………S11 |
| **Text S5 –** Temperature Dependence | …………………………S12 |
| **Text S6 –** Negative Control Comparison | …………………………S14 |
| **Text S7 –** RDase Alignment | …………………………S19 |

#### SUPPLEMENTAL TABLES

| | |
|---|---|
| **Table S1.** Expression plasmids | ……………………………S3 |
| **Table S2.** RDase concentration and purity | ……………………………S5 |
| **Table S3**. Reaction buffer vial setup | ……………………………S7 |
| **Table S4.** Raw data for enzyme assays. | Accompanying Excel |
| **Table S5.** TmrA kinetic values and model statistics | Accompanying Excel |
| **Table S6.** CfrA kinetic values and model statistics | Accompanying Excel |
| **Table S7.** AcdA kinetic values and model statistics | Accompanying Excel |
| **Table S8.** TCE inhibition kinetic values and model statistics | Accompanying Excel |
| **Table S11.** CF kinetic values at varying temperatures | ……………………………S14 |
| **Table S12.** Residues 260, 286, and 319 of each OG 97 member. | ……………………………S19 |

#### SUPPLEMENTAL FIGURES

| | |
|---|---|
| **Figure S1.** SDS-PAGE of RDase purifications | …..………………………S5 |
| **Figure S2.** Schematic of the kinetic assay setup and process | ..…………………………S8 |
| **Figure S3.** Enzyme reaction schemes | …………………………S11 |
| **Figure S4.** CF kinetic curves at varying temperatures | …………………………S13 |
| **Figure S5.** Raw production of DCM in CF assays and negative controls | …………………………S15 |
| **Figure S6.** Conversion of CF to DCM | ……….……………..…S15 |
| **Figure S7.** Raw production of 1,1-DCA in 1,1,1-TCA assays and negative controls | …………………………S16 |
| **Figure S8.** Conversion of 1,1,1-TCA to 1,1-DCA | ……….…..…………S16 |
| **Figure S9.** Raw production of 1,2-DCA and VC in 1,1,2-TCA assays and negative controls | …………..……………S17 |
| **Figure S10.** Conversion of 1,1,2-TCA to 1,2-DCA and VC | …………………………S17 |
| **Figure S11.** Percent of 1,2-DCA and VC produced at varying 1,1,2-TCA concentrations | …………………………S18 |
| **Figure S12.** Raw production of DCM in CF with TCE inhibition assays and negative controls | …………………………S18 |

### Text S1 – RDase Expression and Purification

This section contains methodology for the expression and partial purification of the three RDases TmrA, CfrA, and AcdA, and the concentrations of each enzyme used to calculate the rate of reaction. The construction of the expression plasmids *p15TVL-tmrA*, *p15TVL-cfrA*, and *p15TVL-acdA* and the production of their expression strains have been described elsewhere (Table S1) [1,2]. Each expression plasmid was expressed in *E. coli* BL21(DE3) *cat-araC*-$P_{BAD}$-*suf*, *ΔiscR::kan, ΔhimA::*$Tet^R$ with co-expression of the *pBAD42-BtuCEDFB* plasmid [3,4]. The *pBAD42-BtuCEDFB* plasmid was generously provided by the Booker Lab (Pennsylvania State University, PA, USA). *E. coli* BL21(DE3) *cat-araC*-$P_{BAD}$-*suf*, *ΔiscR::kan, ΔhimA::*$Tet^R$, was generously provided by the Antony and Kiley Labs (St. Louis University School of Medicine, MO, USA; University of Wisconsin-Madison, WI, USA).

**Table S1.** Expression plasmids used in this work.

| Plasmid | Encoded Enzyme | Tags | Induction | Antibiotic Resistance | Source |
|---|---|---|---|---|---|
| *p15TVL-CfrA* | CfrA (no TAT) | N-term 6xHis | IPTG | Ampicillin | [2] |
| *p15TVL-AcdA* | AcdA (no TAT) | N-term 6xHis | IPTG | Ampicillin | [1] |
| *p15TVL-TmrA* | TmrA (no TAT) | N-term 6xHis | IPTG | Ampicillin | [2] |
| *pBAD42-BtuCEDFB* | BtuB, BtuC, BtuD, BtuE, BtuF | N/A | Arabinose | Spectinomycin | Booker lab (Pennsylvania State University) [4] |

**Enzyme Expression**

Large-scale expression of the RDases was carried out in 2 L media bottles with 1 L of Luria-Broth (LB) supplemented with carbenicillin and spectinomycin antibiotics. The 1 L culture was inoculated with 5 mL of overnight culture and grown with a foil cap, aerobically at 37°C, 170 rpm until the $OD_{600}$ reached 0.3-0.5. The culture was then supplemented with a final concentration of 3 µM hydroxycobalamin hydrochloride, and the Btu pathway was induced with 0.2% L-arabinose. The bottle was then capped with a silicon septum, and the culture was further incubated at 37°C until the $OD_{600}$ reached 0.6-0.8. The bottle was then put on ice and the culture was purged with $N_2$ gas for 30 min. The anaerobic culture was supplemented with final concentrations of 50 µM ammonium ferric citrate and 50 µM cysteine from anaerobic 1000x stocks. The RDase production was induced with 0.5 mM isopropyl β-d-1-thiogalactopyranoside (IPTG). The culture was finally incubated overnight (18-20 hr) at 15°C, 170 rpm for protein production. After incubation, in an anaerobic glovebox, the culture was transferred to a 1 L o-ringed centrifuge bottle that had been equilibrated under anaerobic conditions. The culture was pelleted by centrifugation at 4500xg for 20 min. The supernatant was discarded, and the pellet was sealed in the bottle under anaerobic conditions and stored at -80°C until purification. This process was the same for each TmrA, CfrA, and AcdA.

**Enzyme Purification**

All lysis and purification steps were carried out in an anaerobic glovebox in an atmosphere of 10% $CO_2$/10% $H_2$/Bal. $N_2$ (v/v), unless otherwise stated. The pellet was thawed on ice and transferred into the glovebox. The pellet was resuspended in 10 mL of anaerobic lysis buffer [50 mM Tris-HCl pH 7.5, 150 mM NaCl, 0.1% Triton X-100, 5% glycerol, 1 mM tris(2-carboxyethyl)phosphine (TCEP), 50 µg/mL leupeptin, 2 µg/mL aprotinin, 10 mM $MgCl_2$], the cells were lysed using BugBuster® 10x concentrate protein extraction reagent, 300 µg/mL lysozyme, and 1 µg/mL DNase. The lysis was carried out in a sealed 50 mL rounded-bottom centrifuge tube at room temperature, 70 rpm for 20 min. The lysate was clarified by centrifugation at 35 000xg, 4°C, for 20 min.

In the glovebox, a gravity column with 1 mL anaerobic Ni-NTA resin was equilibrated with Wash Buffer (50 mM Tris-HCl pH 7.5, 150 mM NaCl, 30 mM imidazole, 1 mM TCEP). The clarified lysate was loaded onto the resin, and the flow-through was collected and reloaded on the column to allow for greater enzyme capture. The bound protein was washed with 20-30 column volumes (CVs) of Wash Buffer, until protein could no longer be detected in the eluant by Bradford reagent. The RDase was then eluted from the column using 10 CVs of Elution Buffer (50 mM Tris-HCl pH 7.5, 150 mM NaCl, 300 mM imidazole, 1 mM TCEP), and fractions of ~2 mL were manually collected. All fractions that had protein detectable by Bradford reagent were pooled and concentrated in a 10 kDa cutoff filter tube by centrifugation at 3000xg, 4°C for 30 min. A buffer exchange was then performed by diluting the concentrated protein in 15 mL of Storage Buffer (50 mM Tris-HCl pH 7.5, 150 mM NaCl, 2 mM TCEP). The protein was concentrated again by centrifugation. The concentrated protein was aliquoted in 50 µL volumes and stored in liquid $N_2$. A fresh aliquot was used for each kinetic experiment. The protein concentration was measured by Bradford assay using triplicate measurements, and the purity was estimated by sodium dodecylsulfate polyacrylamide gel electrophoresis (SDS-PAGE) using Image Lab 6.1 software (2022, Bio-Rad Laboratories, Inc.), (Figure S1, Table S1).

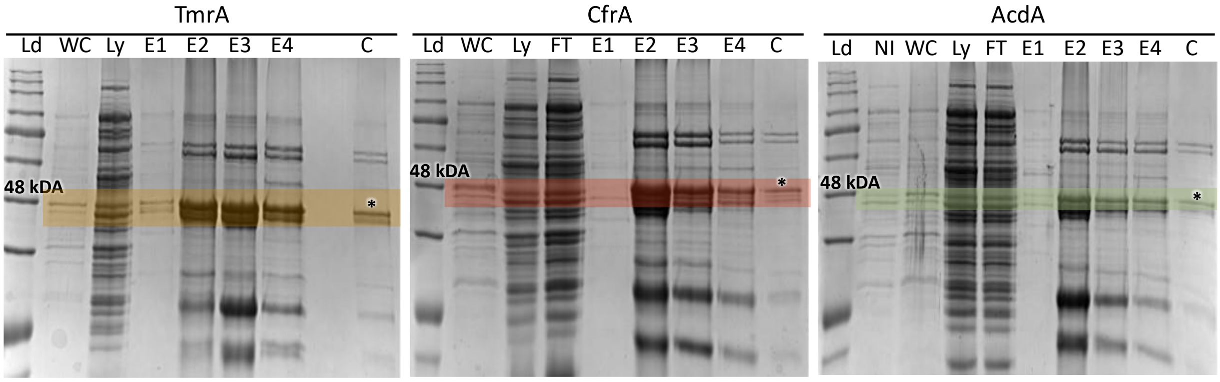

**Figure S1.** SDS-PAGE showing the nickel affinity clean-up of TmrA (46 kDa), CfrA (46 kDa), and AcdA (46 kDa). Ld = FroggaBio BlueEYE pre-stained protein ladder, NI = whole cell not-induced, WC = whole cell post-induction, Ly = soluble lysate fraction, FT = flow through, E = elution fraction, C = combined protein after buffer exchange. The RDase in the combined fraction is indicated by an asterisk.

**Table S2.** Protein concentration for each RDase, the estimated purity from the SDS-PAGE, and the resulting concentration and standard deviation used for rate calculations. Standard deviation is from triplicate Bradford assay measurements.

| Protein | Protein Concentration (mg/mL) | STD (mg/mL) | Estimated Purity | RDase Concentration Accounting for Purity (mg/mL) | STD (mg/mL) |
|---|---|---|---|---|---|
| TmrA | 1.73 | 0.15 | 25% | 0.43 | 0.04 |
| CfrA | 1.86 | 0.22 | 20% | 0.37 | 0.04 |
| AcdA | 1.54 | 0.16 | 20% | 0.31 | 0.03 |

### Text S2 – Detailed Kinetic Assay Methods

This section contains details on the setup of the reaction buffer vials and the procedure of running the assay. The reaction vials were set up with 50 mM Tris-HCl pH 7.5 and 1 mM methyl viologen at the volumes stated in Table S3 (volumes vary such that the final volume with substrate and reductant was 6 mL of liquid), the amount of either saturated solvent stock or pure solvent added to get the desired liquid concentration in the vial was supplemented with Hamilton syringes and this was equilibrated for 24 hr. Immediately before the assays were performed, 5 mM Ti(III) citrate was supplemented using 0.375 mL of 80 mM stock. Some of the substrates had more than 15 substrate levels due to gaps in the Michaelis-Menten curve or outliers; 1,1,1-TCA had additional higher concentrations added because some samples were removed as outliers, 1,1,2-TCA had a few additional low concentrations added because the observed $K_M$

values were lower than initially thought based on literature. The pure solvent was used to deliver the substrate if the volume of the saturated stock was too high or, in the case of 1,1,1-TCA, to try and mitigate the presence of the acetic acid, which is a known hydrolysis product to 1,1,1-TCA. The saturated stocks were assumed to be at the concentration of solubility for each substrate in water. The aqueous concentration in the vial was slightly higher than the target final assay concentration to account for dilution when the enzyme was added. The actual concentration in the vial does not impact the assay very much because the initial concentration is measured based on the remaining substrate and product in a closed vessel so there should be no loss.

For the trichloroethene (TCE) inhibited assay, 50 mL of buffer (with no reductant) was set up in 110 mL bottles with 50 μM TCE added using 790 μL of a TCE saturated water stock (a bottle was also set up with no TCE). This was then used to set up the CF vials as normal, with only ten substrate levels. A schematic of the vials setup and assay procedure is in Figure S2.

**Analytical Procedures (Gas Chromatography**

Gas chromatography (GC) analysis was carried out on an Agilent 7890A GC instrument with flame ionization detection (FID) equipped with an Agilent G1888 headspace autosampler. All samples were measured with the same instrument method using an Agilent GS-Q plot column (30 m length, 0.53 mm diameter). Helium was used as the carrier gas with an 11 mL/min flow rate. The 11-mL headspace autosampler vials containing acidified samples were loaded on the autosampler tray, where they were first equilibrated to 70°C for 40 min, then 3 mL of headspace was sampled via a heated transfer line and injected onto the column by a packed inlet maintained at 200°C. After injection, the oven was held at 35°C for 1.5 min then the temperature was ramped up 15°C/min until 100°C was reached. The ramp rate was then reduced to 5°C/min until 185°C was reached where the temperature was held for 10 min. Finally, the column was cleared by ramping the temp 20°C/min to 200°C and holding for 10 min. The FID detector was set to 250°C and data were collected and analyzed using Agilent ChemStation Rev. B.04.02 SP1 software.

External calibration curves were used for each substrate and product to translate the chromatogram peak area to liquid concentration and total nmol per reaction volume. Standard mixes containing all substrates and products of interest were made in sealed methanolic stocks to prevent partitioning into headspace. Standards are prepared by measuring the desired amount of methanolic stock into HCl acidified water to a final volume of 6 mL in the 11-mL headspace vials and immediately crimped. Standard curves were created using five known liquid concentration levels of the standard mix, each in triplicate. The response factor for each compound, derived from the standard curve, was used to convert peak areas to liquid concentrations. One standard was run with each GC-FID batch to ensure consistent calibration.

**Table S3.** The concentrations of each substrate and the amount of stock used in each reaction buffer stock vial used in the kinetic assays.

| Substrate | Target Liquid Concentration in Syringe (µM) | Target Liquid Concentration in Vial (µM) | Volume of Saturated Stock[a] (µL) | Volume of Pure Solvent (µL) | Buffer volume (mL) |
|---|---|---|---|---|---|
| CF | 0 | 0 | 0 | - | 5.625 |
| CF | 10 | 11.1 | 1.1 | - | 5.624 |
| CF | 20 | 22.2 | 2.2 | - | 5.623 |
| CF | 40 | 44.4 | 4.4 | - | 5.621 |
| CF | 50 | 55.6 | 5.5 | - | 5.620 |
| CF | 60 | 66.7 | 6.6 | - | 5.619 |
| CF | 80 | 88.9 | 8.8 | - | 6.616 |
| CF | 100 | 111 | 11.0 | - | 5.614 |
| CF | 125 | 139 | 13.8 | - | 5.611 |
| CF | 150 | 167 | 16.6 | - | 5.609 |
| CF | 200 | 222 | 22.1 | - | 5.603 |
| CF | 300 | 333 | 33.1 | - | 5.592 |
| CF | 500 | 556 | 55.3 | - | 5.570 |
| CF | 750 | 834 | 82.9 | - | 5.543 |
| CF | 1000 | 1111 | 110 | - | 5.515 |
| CF | 1500 | 1667 | 166 | - | 5.459 |
| 1,1,1-TCA | 0 | 0 | 0 | - | 5.625 |
| 1,1,1-TCA | 5 | 5.5 | 5.4 | - | 5.620 |
| 1,1,1-TCA | 10 | 11.1 | 10.9 | - | 5.614 |
| 1,1,1-TCA | 20 | 22.2 | 21.8 | - | 5.603 |
| 1,1,1-TCA | 40 | 44.4 | 43.7 | - | 5.581 |
| 1,1,1-TCA | 50 | 55.6 | 54.7 | - | 5.570 |
| 1,1,1-TCA | 60 | 66.7 | 65.6 | - | 5.559 |
| 1,1,1-TCA | 80 | 88.9 | 87.5 | - | 5.538 |
| 1,1,1-TCA | 100 | 111 | 109 | - | 5.516 |
| 1,1,1-TCA | 150 | 167 | 164 | - | 5.461 |
| 1,1,1-TCA | 200 | 222 | 219 | - | 5.407 |
| 1,1,1-TCA | 250 | 278 | 273 | - | 5.352 |
| 1,1,1-TCA | 300 | 333 | 328 | - | 5.297 |
| 1,1,1-TCA | 350 | 389 | 383 | - | 5.242 |
| 1,1,1-TCA | 400 | 444 | 437 | - | 5.188 |
| 1,1,1-TCA | 500 | 556 | - | 0.5 | 5.625 |
| 1,1,1-TCA | 750 | 834 | - | 0.8 | 5.624 |
| 1,1,1-TCA | 1000 | 1111 | - | 1.1 | 5.624 |
| 1,1,1-TCA | 1500 | 1667 | - | 1.6 | 5.623 |
| 1,1,2-TCA | 0 | 0 | 0 | - | 5.625 |
| 1,1,2-TCA | 10 | 11.1 | 2.0 | - | 5.623 |
| 1,1,2-TCA | 25 | 27.8 | 5.0 | - | 5.620 |
| 1,1,2-TCA | 50 | 55.6 | 9.9 | - | 5.615 |
| 1,1,2-TCA | 75 | 83.3 | 15 | - | 5.610 |
| 1,1,2-TCA | 100 | 111 | 20 | - | 5.605 |
| 1,1,2-TCA | 250 | 278 | 50 | - | 5.575 |
| 1,1,2-TCA | 500 | 556 | 100 | - | 5.525 |
| 1,1,2-TCA | 750 | 833 | 150 | - | 5.475 |
| 1,1,2-TCA | 1000 | 1110 | 200 | - | 5.425 |
| 1,1,2-TCA | 1250 | 1390 | 250 | - | 5.375 |
| 1,1,2-TCA | 1500 | 1670 | 300 | - | 5.325 |
| 1,1,2-TCA | 1750 | 1940 | - | 1.10 | 5.624 |
| 1,1,2-TCA | 2000 | 2220 | - | 1.27 | 5.624 |
| 1,1,2-TCA | 2500 | 2780 | - | 1.60 | 5.623 |

| | | | | | |
|---|---|---|---|---|---|
| 1,1,2-TCA | 3000 | 3330 | - | 1.91 | 5.623 |
| 1,1,2-TCA | 4000 | 4440 | - | 2.54 | 5.622 |
| 1,1,2-TCA | 5000 | 5560 | - | 3.19 | 5.622 |

[a] assumed concentrations are 67.8 mM CF, 9.7 mM 1,1,1-TCA, and 34.4 mM 1,1,2-TCA
CF = chloroform, TCA = trichloroethane

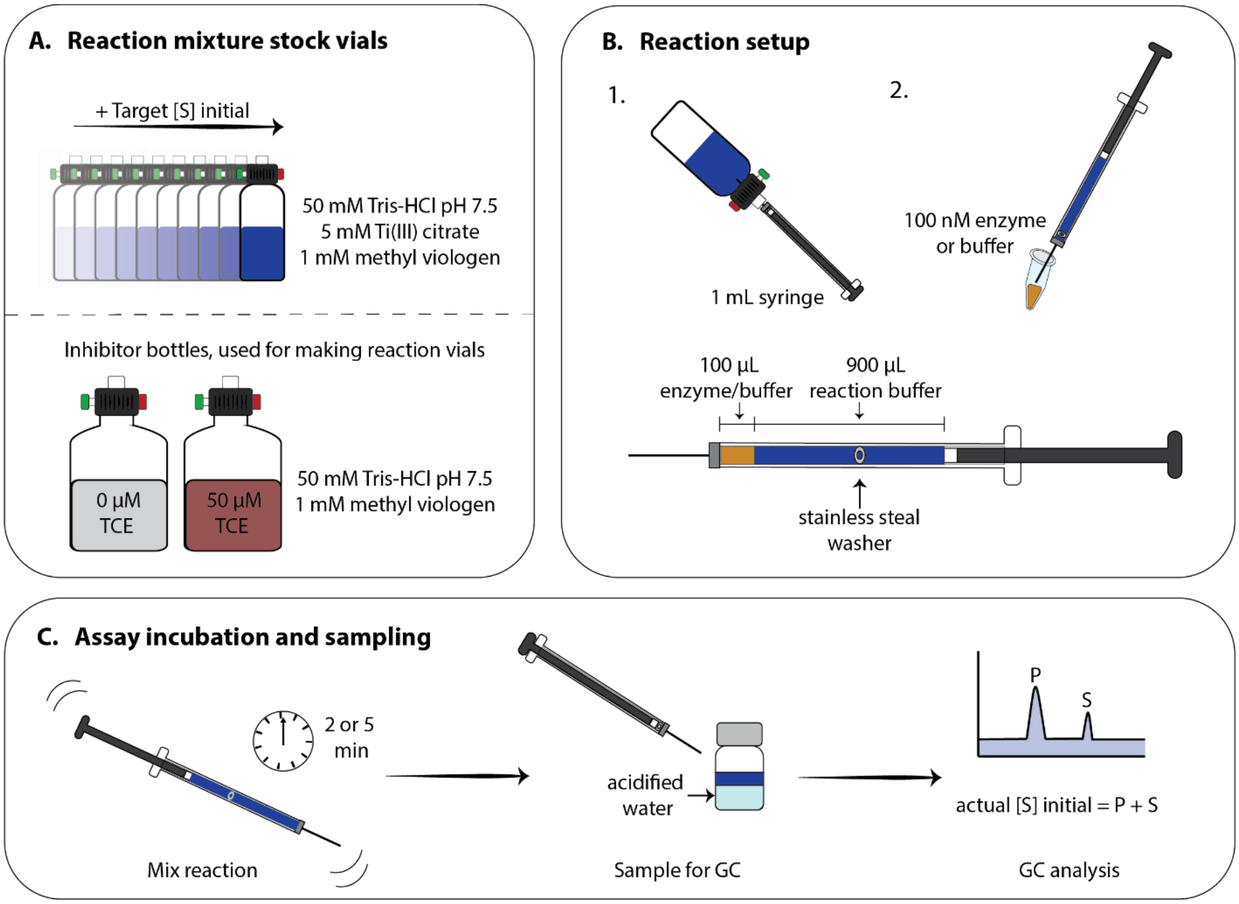

**Figure S2.** Schematic of the kinetic assay setup and process. (A) Depiction of the reaction buffers in vials with increasing target initial substrate concentrations ([S] initial). The two bottom bottles (grey and red) are used in the trichloroethene (TCE) inhibition assay as an initial buffer stock to ensure the individual vials have the same inhibitor concentrations. (B) The reaction setup using a 1 mL glass gas-tight Hamilton syringe; 900 µL of buffer from a reaction vial is taken up first and then 100 µL of the desired enzyme (or buffer) is taken up. (C) The incubation steps with agitation of the syringe and washer, sampling for gas chromatography (GC) and determination of the actual initial substrate concentration from the product (P) and leftover substrate (S) measured by GC.

### Text S3 – Enzyme Kinetic Models

The following equations were used to fit the kinetic data. Equation 1 is the classic Michaelis-Menten model of enzyme kinetics. The reaction rate $v$ (nmol/s or $s^{-1}$) is dependent on the kinetic constants and the substrate concentration [S]. The constants are $V_{max}$ (nmol/s), the maximum rate of catalysis; $K_M$ (µM), the half-saturation co-efficient; $k_{cat}$ ($s^{-1}$), the first-order rate constant; [E], the concentration of the enzyme. Equation 2 gives the relationship between $V_{max}$ and $k_{cat}$. The kinetic constants will vary depending on the enzyme and the substrate involved. These constants are used throughout the rest of the models.

Equation 3 is a model that accounts for inhibition by high concentrations of the substrate. This model has another constant, $K_{I,\,sub}$, the inhibition constant of the substrate. The competitive (Equation 4), uncompetitive (Equation 5), and non-competitive (Equation 6) models are used in a reaction where there is an external inhibitor added; the fit of the model will depend on the mode of action of the inhibitor. In these models [I] is the concentration of the inhibitor, and $K_{I,inh}$ is the inhibition constant of the inhibitor. These inhibition models can be combined with substrate inhibition to give Equations 7-9.

Competitive inhibition is when the external inhibitor binds to the substrate-binding pocket and blocks the substrate from binding; this will only affect the apparent $K_M$ because excess amounts of substrate can outcompete the inhibitor. In uncompetitive inhibition, the inhibitor binds to the enzyme-substrate complex rendering the enzyme inactive; since the same amount of substrate binds the enzyme but less is transformed, this will only affect the apparent $V_{max}$. Non-competitive inhibition is a combination: the inhibitor can bind to the free enzyme and the enzyme-substrate complex causing an effect on both the apparent $K_M$ and $V_{max}$.

Equation 1- Michaelis-Menten Model
$$v = \frac{V_{max}[S]}{K_M+[S]} = \frac{k_{cat}[E][S]}{K_M+[S]}$$

Equation 2- $V_{max}$ and $k_{cat}$ relationship
$$V_{max} = k_{cat}[E]$$

Equation 3- Substrate Inhibition Model
$$v = \frac{V_{max}[S]}{K_M+[S]\left(1+\frac{[S]}{K_{I_{sub}}}\right)}$$

Equation 4- Competitive Inhibitor Model
$$v = \frac{V_{max}[S]}{K_M\left(1+\frac{[I]}{K_{I_{inh}}}\right)+[S]}$$

Equation 5- Uncompetitive Inhibitor Model
$$v = \frac{V_{max}[S]}{K_M+[S]\left(1+\frac{[I]}{K_{I_{inh}}}\right)}$$

Equation 6- Non-competitive Inhibitor Model

$$v = \frac{V_{max}[S]}{K_M\left(1+\frac{[I]}{K_{I_{inh}}}\right)+[S]\left(1+\frac{[I]}{K_{I_{inh}}}\right)}$$

Equation 7- Competitive Inhibitor with Substrate Inhibition

$$v = \frac{V_{max}[S]}{K_M\left(1+\frac{[I]}{K_{I_{inh}}}\right)+[S]\left(1+\frac{[S]}{K_{I_{sub}}}\right)}$$

Equation 8- Uncompetitive Inhibitor with Substrate Inhibition

$$v = \frac{V_{max}[S]}{K_M+[S]\left(1+\frac{[I]}{K_{I_{inh}}}\right)\left(1+\frac{[S]}{K_{I_{sub}}}\right)}$$

Equation 9- Non-competitive Inhibitor with Substrate Inhibition

$$v = \frac{V_{max}[S]}{K_M\left(1+\frac{[I]}{K_{I_{inh}}}\right)+[S]\left(1+\frac{[I]}{K_{I_{inh}}}\right)\left(1+\frac{[S]}{K_{I_{sub}}}\right)}$$

In Michaelis-Menten kinetics, the overall reaction has two general steps: the reversible formation of the enzyme-substrate complex (ES) and the reaction of the ES to release the product (Figure S3A). The first-order rate of product formation is, therefore, dependent on the kinetic coefficient ($k_2$) and the concentration of the ES. This is transformed to Equation 1 using the steady-state assumption and defining the kinetic parameter $K_M$ as the dissociation constant that accounts for the equilibrium to form ES and the consumption to release the product. In the case of 1,1,2-TCA there are two possible products, 1,2-DCA and VC. Each enzyme forms a different mixture of products and, as such, most likely have different rates of product formation. To measure these rates, we need to look at a new reaction scheme where the ES can take two paths each with its own rate coefficient (Figure S3B). The $K_M$ will account for both rates of product formation meaning that the reaction rate for each product will depend on each other. To account for this relationship, we first modelled the overall consumption of 1,1,2-TCA with the sum of the products to get the $K_M$ and $K_{I,sub}$ values. Then, the products were separated to be modelled again with these values fixed in the model in order to obtain the $V_{max}$, which will only account for the ES to product transformation step.

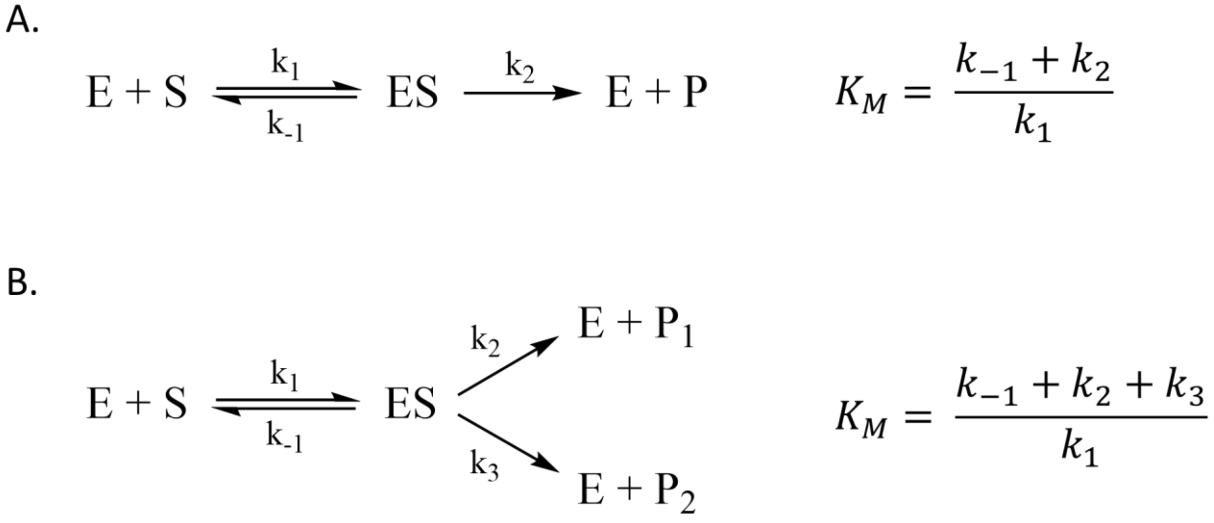

**Figure S3.** Enzyme reaction schemes and $K_M$ values for (A) the typical one-product reaction and (B) the two-product reaction.

### Text S4 – Raw Data and Model Fits

All raw data for the assays performed are in Table S4A-D, provided in the accompanying Excel file. The raw data includes the temperature of each reaction, the mass of the substrate and product measured and the target and actual initial concentration of the substrate. The target concentration varies from the actual concentration due to discrepancies in the saturated stock solution used to deliver most of the substrates (concentration assumed to be the solubility of each compound in water), error in the small volumes of pure solvent added for some mid-range or high concentrations, and changes to the liquid concentration as buffer was taken and headspace was introduced. The outliers removed for 1,1,1-TCA analysis are also identified.

The enzyme kinetics on CF were compared with no TCE and with ~45 µM TCE in the assay mixture. The $K_{I,TCE}$ of TCE was calculated using the same $V_{max}$, $K_M$, and $K_{I,CF}$ (TmrA and AcdA only) calculated from the uninhibited data taken on the same day. All enzymes were fit with competitive, uncompetitive, and non-competitive inhibition models; the best-fit model was chosen as the mode of inhibition. TmrA and AcdA had the best fit with uncompetitive type inhibition, whereas CfrA had the best fit with non-competitive inhibition. However, a full inhibition series to determine that CfrA demonstrated uncompetitive inhibition by TCE [5].

The assessment and results for the class Michaelis-Menten and substrate inhibition models for TmrA, CfrA, and AcdA kinetics are shown in Tables S5, S6, and S7, respectively. The TCE inhibition experiments were fit with competitive, non-competitive, and uncompetitive kinetic models, the assessment of the models for each enzyme is in Table S8. The kinetic models were compared using the fit $R^2$, the Akaike information criterion (AIC), and the standard deviation of the residuals ($S_{x,y}$). Generally, a good model fit will have an $R^2$ value close to 1, though this is not the best for comparing model fits. The AIC is a statistical method that evaluates the probability of observing the data under the model and the number of parameters, with a penalty for more parameters to prevent overfitting. The AIC is only used to compare the relative fit of two models on the same dataset, and the model with a lower score is the preferred model. Similarly, $S_{x,y}$ can be used to compare fits of the same data and lower values represent a better model because the observed data is closer to the predicted values. The models that had lower AIC and $S_{x,y}$ values for each data set were used to determine the kinetic parameters.

The uncertainty of each kinetic parameter was calculated from the standard error multiplied by 1.96—the number of standard deviations —to get the 95% confidence interval.

### Text S5 – Temperature Dependence

The reactions were unable to be held at a consistent temperature due to the anaerobic setup. Consequently, repeated experiments had slightly different kinetic rates. This section compares the kinetics of each enzyme using CF as a substrate at two different temperatures.

**Temperature Effect**

A major limitation of the assay setup is the inability to control the temperature—an important factor in enzyme rate. Due to the anaerobic setup, the short incubation time, and the necessity to manually agitate the reaction vessel (i.e. the syringe), the temperature was recorded but could not be altered. We did see a large difference in the $V_{max}$ using CF as a substrate when the temperature varied between 20°C and 24°C (Figure S4, Table S11). The optimal temperature for TmrA is 45°C [6], so it is unsurprising that there is a jump in activity with the higher temperature. TmrA and CfrA's $K_M$ values remained consistent, but AcdA had some variation in the $K_M$ and $K_{I,CF}$ it experienced, with a lower $K_M$ and higher $K_I$ at 20°C. This change in the affinity for CF suggests that the enzyme experiences less inhibition at lower temperatures; possibly the enzyme has a more rigid structure to hold CF in place. The data for CfrA at 20°C is used from the TCE inhibition test, while TmrA and AcdA were done in a full 20°C trial. The full trial of CfrA did not show as drastic a change in the $V_{max}$ as it did in the TCE trial, which may be due to temperature fluctuations over the course of the experiment (e.g. increased temperature of glovebox due to direct sunlight). Data for all CfrA trials are in All raw data for the assays performed are in Table S4C and kinetic model information for all enzymes at the two temperatures is in Tables S9-10.

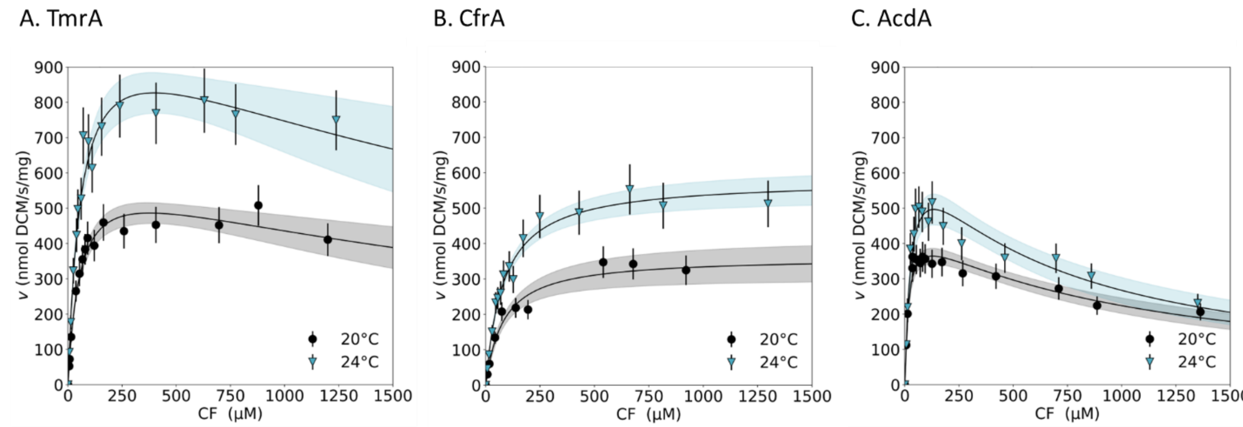

**Figure S4.** Reaction rate against substrate concentration plots for TmrA (A), CfrA (B), and AcdA (B) using chloroform (CF). The reactions were performed at either 20°C (black circles) or 24°C (blue triangles). Error bars on the plot represent the uncertainty in the protein concentration and incubation time propagated in the calculated rate. The black lines are the highest-scoring model kinetic model for each set of data. The shaded area around the solid lines represents the uncertainty in the predicted model parameters. Kinetic assays were performed with a reaction mixture of 10 nM RDase, 5 mM Ti(III) citrate, 1 mM methyl viologen in 50 mM Tris-HCl pH 7.5 and varying concentrations of the substrate. All reactions were carried out over 2 min.

The activation energy ($E_a$) of the reaction catalyzed by each enzyme was estimated using the Arrhenius equation (Equation 10). The Arrhenius equation gives a relationship between the temperature and the rate of the reaction, and a rule of thumb is that the reaction rate will double for every 10°C increase in temperature because many enzyme-catalyzed reactions have $E_a$ between 40-50 kJ/mol [7]. Enzyme kinetics follow this relationship while the enzyme is active, but enzymes will denature once they reach temperatures where they are no longer stable. Given that the temperature increases by 4°C, we expect that the reaction rate will increase by approximately 1.4x. The calculated maximum rates for these enzymes increase 1.5-1.7x with the temperature increase, the $E_a$ is also a larger number than expected. This larger-than-expected increase may be due to inconsistency between the temperature glovebox atmosphere and the temperature of the reaction liquid. As these are being manually mixed, heat could be transferred to the relatively small reaction volume to increase the temperature unintentionally.

Equation 10 – Arrhenius Equation $$\ln\left(\frac{k_2}{k_1}\right) = \frac{E_a}{R}\left(\frac{1}{T_1} - \frac{1}{T_2}\right)$$

**Table S11.** Kinetic parameters measured for CfrA, TmrA, and AcdA with chloroform as a substrate with incubation temperatures of either 20 or 24°C, and the estimated activation energy ($E_a$).

| | Temperature of Assay | $K_M$ (µM) | $K_{I,CF}$ (µM) | $V_{max}$ (nmol/s/mg) | $k_{cat}$ ($s^{-1}$) | Estimated Activation Energy (kJ/mol) |
|---|---|---|---|---|---|---|
| CfrA | 24°C | 82 ± 12 | n.o. | 580 ± 40 | 28 ± 2 | 90 |
| | 20°C | 81 ± 20 | n.o. | 360 ± 50 | 17 ± 2 | |
| TmrA | 24°C | 61 ± 14 | 2600 ± 1900 | 1100 ± 160 | 53 ± 8 | 100 |
| | 20°C | 55 ± 11 | 2600 ± 1600 | 630 ± 70 | 30 ± 3 | |
| AcdA | 24°C | 29 ± 8 | 610 ± 240 | 710 ± 120 | 34 ± 6 | 80 |
| | 20°C | 17 ± 8 | 960 ± 270 | 460 ± 50 | 22 ± 2 | |

### Text S6 – Negative Control Comparison

This section contains figures of the raw product formation of each kinetic assay series (product mass vs initial substrate concentration) for all three enzymes and for the enzyme free controls in Figure S5, Figure S7, and Figure S9. An enzyme-free control reaction was done for every substrate concentration in each assay, except for the TCE inhibition assays; in these assays, only the substrate-free and the highest concentration of substrate were tested in the enzyme-free control (Figure S12).

The ratio of leftover substrate to substrate converted to product is also shown in Figure S6, Figure S8, and Figure S10. All enzymes have a high conversion of CF to DCM; AcdA obtains almost complete conversion in the low concentrations of CF. Whereas the enzymes have a very low turnover of 1,1,1-TCA, reflective of their high $K_M$ values with this substrate. Finally, AcdA and TmrA have fairly high conversion percentages of 1,12-TCA at low concentrations with a mixture of products, but CfrA has low conversion.

The percentage of each product of 1,1,2-TCA for each reaction in the substrate range is shown in Figure S11. The ratio of 1,2-DCA:VC is very consistent for each enzyme as the amount of 1,1,2-TCA increases.

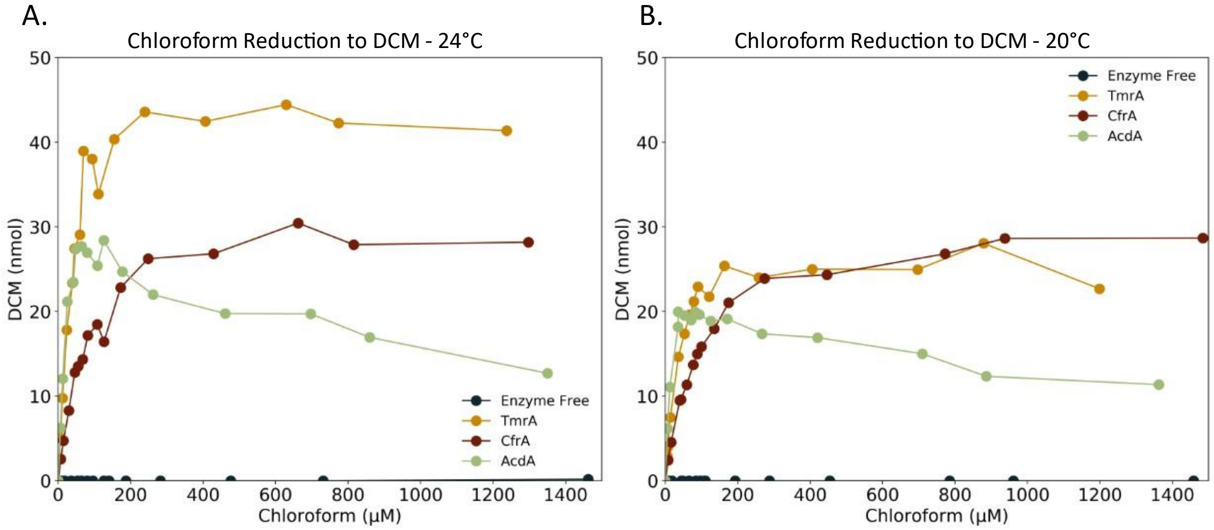

**Figure S5.** Production of dichloromethane (DCM) against chloroform (CF) concentration for reactions incubated with TmrA (yellow), CfrA (red), AcdA (green), or an enzyme-free control (dark teal). (A) is raw data for reactions held at 24°C and (B) is raw data for reactions held at 20°C. Kinetic assays were performed with a reaction mixture of 10 nM RDase, 5 mM Ti(III) citrate, 1 mM methyl viologen in 50 mM Tris-HCl pH 7.5 and varying concentrations of the substrate. All reactions were carried out over 2 min.

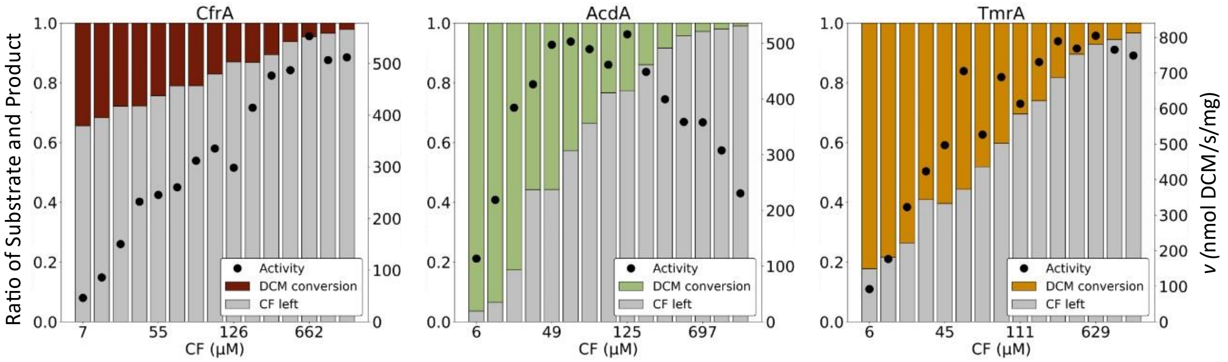

**Figure S6.** Conversion of chloroform (CF) to dichloromethane (DCM) by CfrA (left, red), AcdA (center, green), and TmrA (right, yellow) at different substrate concentrations. Each plot shows the ratio of the unreacted substrate in grey to the amount of substrate converted to DCM in colour. The rate of reaction is shown on the secondary y-axis (black circles). The x-axis has increasing substrate concentration but not on a linear scale; each bar represents one assay vial; exact values and conversion percentage can be seen in Table S4.

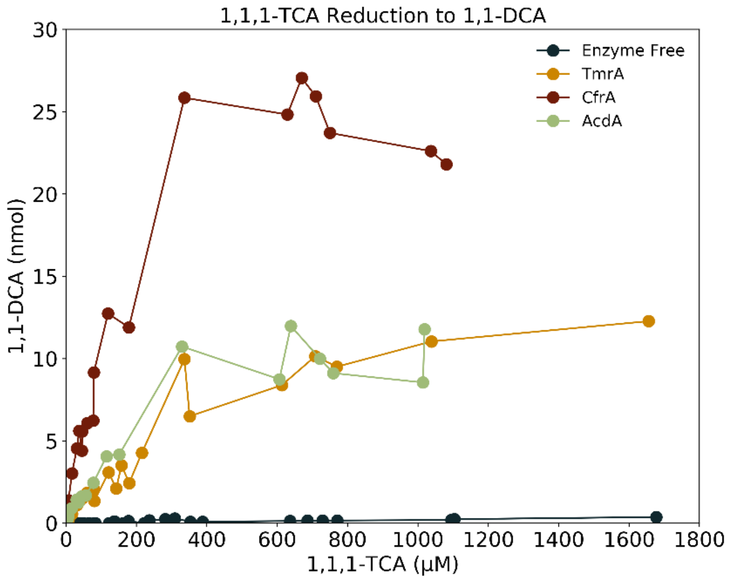

**Figure S7.** Production of 1,1-dichloroethane (1,1-DCA) against 1,1,1-trichloroethane (1,1,1-TCA) concentration for reactions incubated with TmrA (yellow), CfrA (red), AcdA (green), or an enzyme-free control (dark teal). Kinetic assays were performed with a reaction mixture of 10 nM RDase, 5 mM Ti(III) citrate, 1 mM methyl viologen in 50 mM Tris-HCl pH 7.5 and varying concentrations of the substrate. All reactions were carried out over 2 min at 24°C.

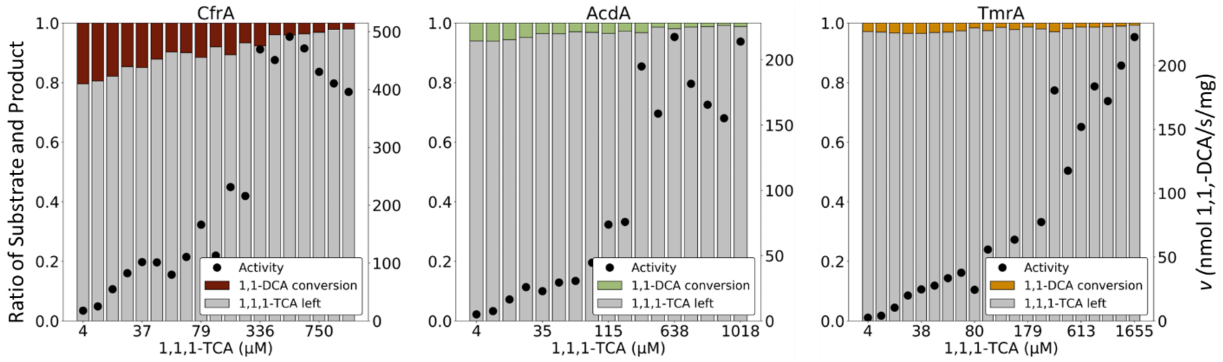

**Figure S8.** Conversion of 1,1,1-trichloroethane (TCA) to 1,1-dichloroethane (DCA) by CfrA (left, red), AcdA (center, green), and TmrA (right, yellow) at different substrate concentrations. Each plot shows the ratio of the unreacted substrate in grey to the amount of substrate converted to 1,1-DCA in colour. The rate of reaction is shown on the secondary y-axis (black circles). The x-axis has increasing substrate concentration but not on a linear scale; each bar represents one assay vial; exact values and conversion percentage can be seen in Table S4.

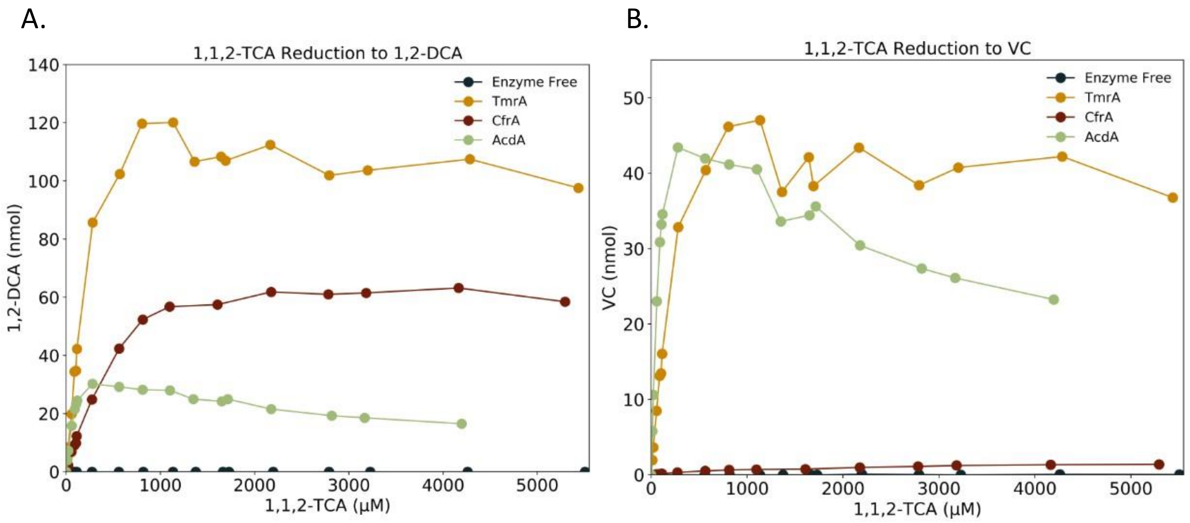

**Figure S9.** Production of (A) 1,2-dichloroethane (1,2-DCA) and (B) vinyl chloride (VC) against 1,1,2-trichloroethane (1,1,2-TCA) concentration for reactions incubated with TmrA (yellow), CfrA (red), AcdA (green), or an enzyme-free control (dark teal). Kinetic assays were performed with a reaction mixture of 10 nM RDase, 5 mM Ti(III) citrate, 1 mM methyl viologen in 50 mM Tris-HCl pH 7.5 and varying concentrations of the substrate. All reactions were carried out over 2 min at 24°C.

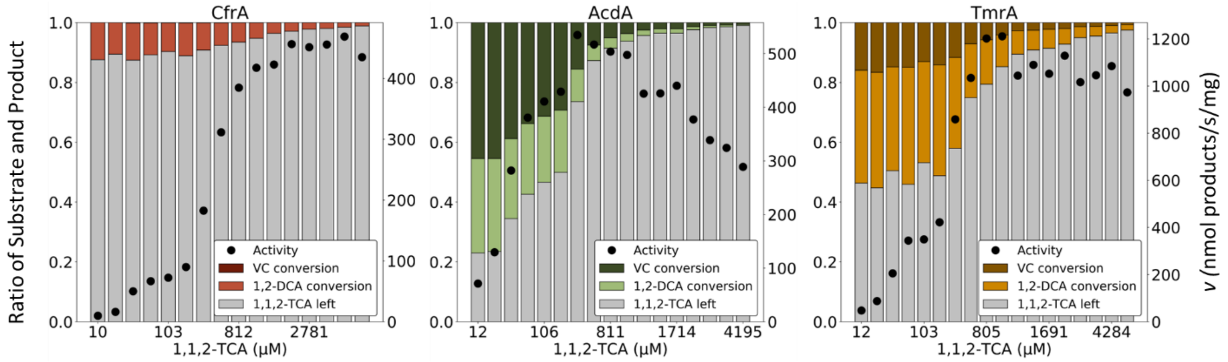

**Figure S10.** Conversion of 1,1,2-trichloroethane (TCA) to 1,2-dichloroethane (DCA) and vinyl chloride (VC) by CfrA (left, red), AcdA (center, green), and TmrA (right, yellow) at different substrate concentrations. Each plot shows the ratio of the unreacted substrate in grey to the amount of substrate converted to 1,2-DCA in the lighter shade and VC in the darker shade of colour. The rate of reaction is shown on the secondary y-axis (black circles). The x-axis has increasing substrate concentration but not on a linear scale; each bar represents one assay vial; exact values and conversion percentage can be seen in Table S4.

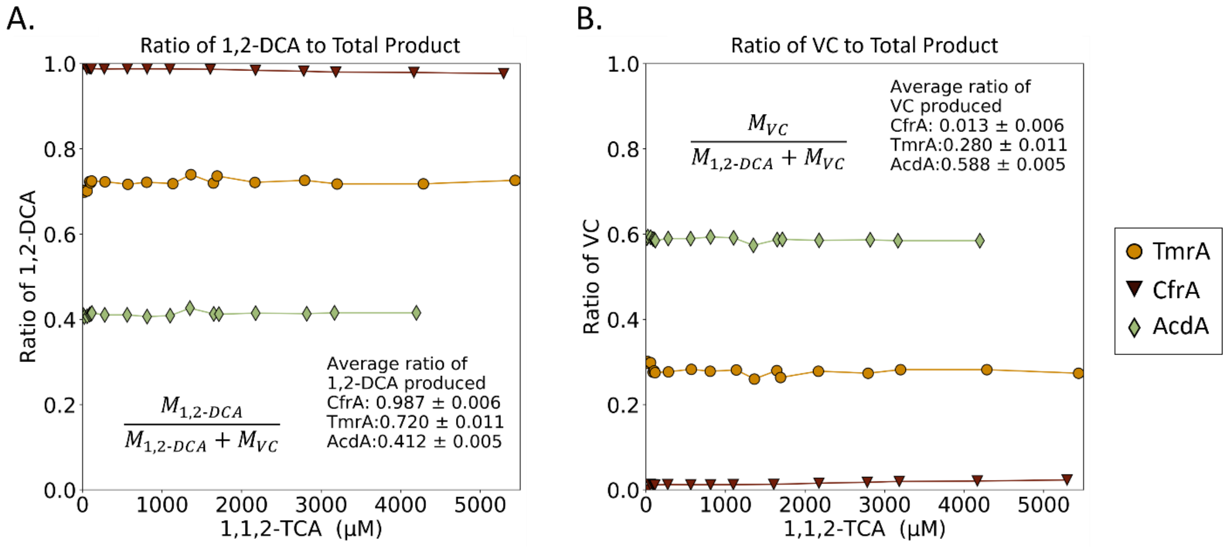

**Figure S11.** The ratio of 1,2-dichloroethane (DCA; A) and vinyl chloride (VC; B) products to the total products formed in the reduction of 1,1,2-trichloroethane by CfrA (red triangles), AcdA (green diamonds), and TmrA (yellow circles) at different substrate concentrations. The equation for calculating the ratio for each point and the average ratio and standard deviation from all substrate concentrations are shown for each enzyme on the graphs.

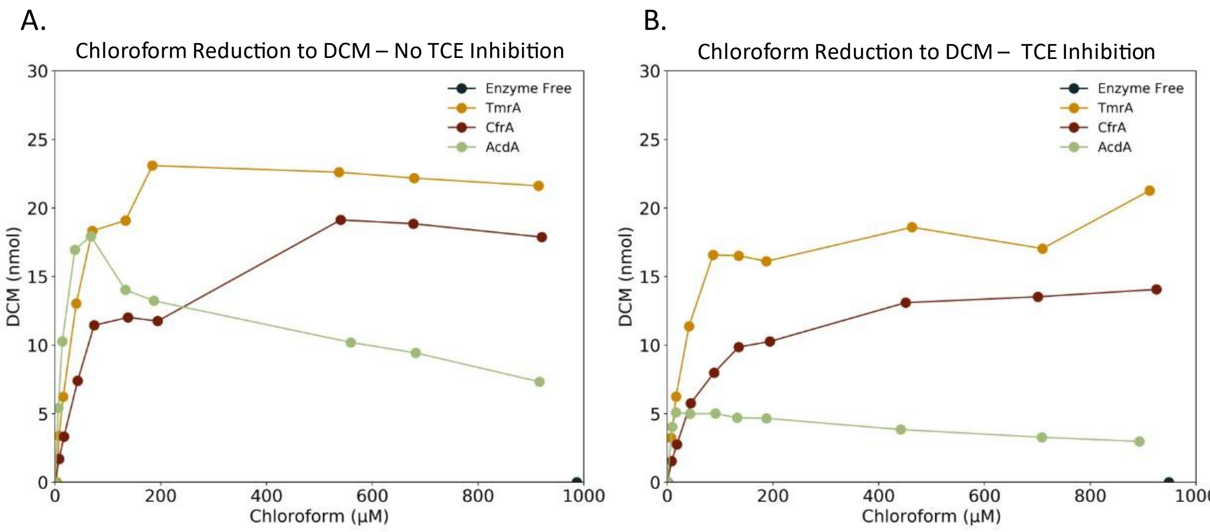

**Figure S12.** Production of dichloromethane (DCM) against chloroform (CF) concentration for reactions incubated with TmrA (yellow), CfrA (red), AcdA (green), or an enzyme-free control (dark teal). (A) is raw data for reactions with no inhibition and (B) is raw data for reactions incubated with trichloroethene (TCE) as an inhibitor. Kinetic assays were performed with a reaction mixture of 10 nM RDase, 5 mM Ti(III) citrate, 1 mM methyl viologen in 50 mM Tris-HCl pH 7.5 and varying concentrations of the substrate. All reactions were carried out over 2 min at 20°C.

### Text S7 – RDase Alignment

This section contains the full amino acid alignments of the three enzymes of focus in this study—TmrA, CfrA, and AcdA—and a selection of characterized RDases from *Dehalococcoides* and *Dehalobacter* to show the three conserved amino acids in highly divergent sequences. Multiple sequence alignments were done using the MUSCLE plugin in Geneious v8.1.9 with default settings.

In the selected sequences, Tyr246 (in PceA) is highly conserved except for TmrA and CfrA. Residue Asn272 in PceA is not as highly conserved with some variation in the *Dehalococcoides* enzymes TceA, VcrA, and BvcA. These enzymes do not conserve the H-bond donor function of Asn727 suggesting that this residue may not be important in all RDases. The conserved arginine (Arg305 in PceA) is highly conserved as either an arginine or lysine, which can perform the same essential proton-donor function.

The putative catalytic residues at positions 260, 286, and 319 for each characterized member of ortholog group 97 are shown in Table S12. Residue Arg319 is conserved in all members, aligning with the hypothesis as a catalytic proton donor. Residues 260 and 286 are variable, with residue 286 seen as either an alanine or a serine and residue 260 as a cysteine, a phenylalanine, or a tyrosine. The variation in residue 260 seems to correlate with each enzyme's 1,1,2-TCA dechlorination product distribution. The enzymes which hold the aromatic residue Phe260 or Tyr260 produce appreciable amounts of VC, whereas CfrA and CtrA hold Cys260 only produce trace amounts of VC. ThmA has not displayed any reduction of 1,1,2-TCA, which could be due to variations elsewhere in the protein structure. The notable difference between the aromatic 260 and cysteine residues suggests that this position could influence either substrate binding or the transformation steps of the reaction. However, we cannot ascertain that this is the sole residue involved because the RDases that have the same residues shown in Table S12 generally have higher overall amino acid sequence identity to each other compared to other OG 97 members. Further, there is not an obvious correlation of these residues of interest in activity on 1,1,1-TCA and CF and it is likely that surrounding active site residues also influence substrate binding.

**Table S12.** Residues 260, 286, and 319 of each OG 97 member.

| OG 97 RDase | Residue 260 | Residue 286 | Residue 319 | 1,1,2-TCA Products | Ref |
|---|---|---|---|---|---|
| TmrA | F | S | R | 2.6 1,2-DCA:1 VC | This study |
| RdhA D8M_v2_40029 | F | S | R | 2.3 1,2-DCA:1 VC | Soder-Walz, 2022 [8] |
| CfrA | C | S | R | Majority 1,2-DCA | This study |
| CtrA | C | A | R | Majority 1,2-DCA | Zhao, 2015 [9] |
| ThmA | C | A | R | Does not reduce 1,1,2-TCA | Zhao, 2017 [10] |
| AcdA | Y | A | R | 0.7 1,2-DCA: 1 VC | This study |
| DcrA | Y | A | R | Majority VC | Picott, 2022 [2] |

OG = ortholog group, TCA = trichloroethane, DCA = dichloroethane, VC = vinyl chloride

> Amino acid sequence alignment of CfrA, AcdA, and TmrA. The TAT sequence that was removed for expression is underlined. Residues that are not conserved between all three enzymes are bolded. Residues in positions 260, 286, and 319, which align with the conserved PceA catalytic residues, are in red.

CfrA/AcdA Identity: 95.8% (437/456)
CfrA/TmrA Identity: 95.4% (435/456)
AcdA/TmrA Identity: 96.7% (441/456)

| | | | |
|---|---|---|---|
| CfrA | 1 | MDKEKSNNDKPATKINRRQFLKFGAGASSGIAIATAATALGGKSLIDPKQVYAGTVKELD | 60 |
| AcdA | 1 | MDKEKSNNDKPATKINRRQFLKFGAGASSGIAIATAATALGGKSLIDPKQVYAGTVKELD | 60 |
| TmrA | 1 | MDKEKSNNDKPATKINRRQFLKFGAGASSGIAIATAATALGGKSLIDPKQVYAGTVKELD | 60 |
| CfrA | 61 | ELPFNIPADYKPFTNQRNI**Y**GQAVLGVPEPLAL**V**ERFDEVRWNGWQTDGSPGLTVLDGAA | 120 |
| AcdA | 61 | ELPFNIPADYKPFTNQRNI**F**GQAVLGVPEPLAL**Q**ERFDEVRWNGWQTDGSPGLTVLDGAA | 120 |
| TmrA | 61 | ELPFNIPADYKPFTNQRNI**F**GQAVLGVPEPLAL**V**ERFDEVRWNGWQTDGSPGLTVLDGAA | 120 |
| CfrA | 121 | ARASFAVDYY**F**NGENSACRANKGFFEWHPKV**A**ELNFKWGDPERNIHSPGVKSAEEGTMAV | 180 |
| AcdA | 121 | ARASFAVDYY**L**NGENSACRANKGFFEWHPKV**P**ELNFKWGDPERNIHSPGVKSAEEGTMAV | 180 |
| TmrA | 121 | ARASFAVDYY**F**NGENSACRANKGFFEWHPKV**P**ELNFKWGDPERNIHSPGVKSAEEGTMAV | 180 |
| CfrA | 181 | K**KI**ARFFGAAKAGIAPFDKRWVFTET**Y**AFVKTPEGE**S**LKFIPPDFGFEPKHVISMIIPQS | 240 |
| AcdA | 181 | K**KI**ARFFGAAKAGIAPFDKRWVFTET**A**AFVKTPEGE**S**LKFIPPDFGFEPKHVISMIIPQS | 240 |
| TmrA | 181 | K**RM**ARFFGAAKAGIAPFDKRWVFTET**A**AFVKTPEGE**D**LKFIPPDFGFEPKHVISMIIPQS | 240 |
| CfrA | 241 | **P**EG**V**KC**D**PSFLGS**T**EYGLS**C**AQIGYAAFGLSMFIKDLGYHAVPIG**S**DSAL**AI**PIAIQAGL | 300 |
| AcdA | 241 | **L**EG**I**KC**A**PSFLGS**A**EYGLS**Y**AQIGYAAFGLSMFIKDLGYHAVPIG**A**DSAL**AV**PIAIQAGL | 300 |
| TmrA | 241 | **L**EG**V**KC**A**PSFLGS**A**EYGLS**F**AQIGYAAFGLSMFIKDLGYHAVPIG**S**DSAL**SI**PIAIQAGL | 300 |
| CfrA | 301 | GEYSRSG**L**MITPEFG**S**NVRLCEVFTDMPLNHDKPISFGVTEFCKTCKKCAEAC**A**PQAISY | 360 |
| AcdA | 301 | GEYSRSG**L**MITPEFG**S**NVRLCEVFTDMPLNHDKPISFGVTEFCKTCKKCAEAC**P**PQAISY | 360 |
| TmrA | 301 | GEYSRSG**Q**MITPEFG**P**NVRLCEVFTDMPLNHDKPISFGVTEFCKTCKKCAEAC**P**PQAISY | 360 |
| CfrA | 361 | EDPTIDGPRGQM**Q**NSGIKRWYVDPVKC**L**EF**M**SRDNV**G**NCCGACIAACPFTKPEAWHHTLI | 420 |
| AcdA | 361 | EDPTIDGPRGQM**H**NSGIKRWYVDPVKC**F**EF**W**SRDNV**R**NCCGACIAACPFTKPEAWHHTLI | 420 |
| TmrA | 361 | EDPTIDGPRGQM**H**NSGIKRWYVDPVKC**F**EF**W**SRDNV**R**NCCGACIAACPFTKPEAWHHTLI | 420 |
| CfrA | 421 | RSLVGAPVITPFMKD**M**DDIFGYGK**L**N**D**EKA**I**ADWWK | 456 |
| AcdA | 421 | RSLVGAPVITPFMKD**V**DDIFGYGK**P**N**D**EKA**I**ADWWK | 456 |
| TmrA | 421 | RSLVGAPVITPFMKD**M**DDIFGYGK**P**N**-**EKA**K**ADWWK | 455 |

> Amino acid sequence alignment of:
*Nitratireductor pacificus* pH-3B NpRdhA (Accession: EKF18105),
*Sulfurospirillum multivorans* PceA (AHJ12791),
*Dehalococcoides mccartyi* PceA (AAW40342),
*Dehalobacter restrictus* PceA (AHF10727),
*Dehalococcoides myccarti* TceA (AAW39060),
*Dehalococcoides mccartyi* VcrA (WP_012882535),
*Dehalococcoides mccartyi* BvcA (AAT64888),
*Dehalobacter* sp. TeCB1 TcbA(WP_068882928),
*Dehalobacter* sp. UNSWDHB TmrA (WP_034377773),
*Dehalobacter* sp. CF CfrA (AFV05253),
*Dehalobacter* sp. SC05-UT AcdA (XCH81453).

Residues in positions which align with the conserved *Sulfurospirillum multivorans* PceA catalytic residues—Tyr246, Asn272, Arg305—are in red. Brackets indicate the organism for enzymes with the same name, Sfs = *Sulfurospirillum*, Dcc = *Dehalococcoides*, Dhb = *Dehalobacter*.

```
NpRdhA        1    -MRLYSNRDRPNHLGPLALERLARVDDVVAQPARQPEDGFAA--------SEDS-LLGDV
PceA (Sfs)    1    ------MEKKKK--PELSRRDFGKLIIGGGAAVTIAPFGVPG--------ANAAEKEKNA
PceA (Dcc)    1    -------MLNFH--STLTRKDFLKGIGMAGAGLGAASAVAPM-------FHDLDELVAST
PceA (Dhb)    1    -----------M--GEINRRNFLKASMLGAAAAAVASASAVK--------GMVSPLVADA
TceA          1    ------MSEKYH--STVTRRDFMKRLGLAGAGAGALGAAVLAENNLPHEFKDVDDLLSAG
VcrA          1    -------MSKFH--KTISRRDFMKGLGLAGAGIGAVAASAPV-------FHDIDELVSSE
BvcA          1    -------MHNFH--CTISRRDFMKGLGLAGAGIGAATSVMPN-------FHDLDEVISAA
TcbA          1    -----------M--GEINRRNFLKASMLGAAAAAVASASAVK--------GMVSPLVADA
TmrA          1    MDKEKSNNDKPA--TKINRRQFLKFGAGASSGIAIATAATAL--------GGKS-LIDPK
CfrA          1    MDKEKSNNDKPA--TKINRRQFLKFGAGASSGIAIATAATAL--------GGKS-LIDPK
AcdA          1    MDKEKSNNDKPA--TKINRRQFLKFGAGASSGIAIATAATAL--------GGKS-LIDPK

NpRdhA        61   EEYARLFTR--FLDGPVAPLGDAIPDDPARRA-----ENLKASAYFLDASMVGICRLDPD
PceA (Sfs)    61   AEIRQQFAM--TAGSPIIVNDKLERYAEVRTAFTHPTSFFKPNYKGEVKPWFLSA-YDEK
PceA (Dcc)    61   PS---------TRNLPW--------FVKER-------EHGDPTT---PIDWDMI-QRRPY
PceA (Dhb)    61   ADIVAPITE--TSEFPYKVDAKYQRYNSLK-------NFFEKTFDPEANKTPIKFHYDDV
TceA          61   KALEGDHANK-VNNHPW--------WVTTR-------DHEDPTC---NIDWSLIKRYSGW
VcrA          61   ANS--------TKDQPW--------YVKHR-------EHFDPTI---TVDWDIFDRYDGY
BvcA          61   SAETSSLSGKSLNNFPW--------YVKER-------DFENPTI---DIDWSILARNDGY
TcbA          61   ADIVAPITE--TSEFPYKVDAKYQRFNCLK-------NFYEKAFDPEANKTPIKFHYDDV
TmrA          61   QVYAGTVKE--LDELPFNIPADYKPFTNQR-------NIFGQAVLGVPEPLALVERFDEV
CfrA          61   QVYAGTVKE--LDELPFNIPADYKPFTNQR-------NIYGQAVLGVPEPLALVERFDEV
AcdA          61   QVYAGTVKE--LDELPFNIPADYKPFTNQR-------NIFGQAVLGVPEPLALQERFDEV

NpRdhA        121  DRAGDCDPSHTHALVFAVQFGREPEAGEAGAEWIRGTNAARTDMRCAEIAAILSGYVRWM
PceA (Sfs)    121  VRQI--------------------------------------------------------
PceA (Dcc)    121  TW-------------------------------VRMDPTLPVYDNLKSIGAPVSRWLDWE
PceA (Dhb)    121  SKITGKKDT-----------GKDLPTLNAERLGIKGRPATHTETSIL-FHTQHLGAMLTQ
TceA          121  NNQGAYF--LPED-------YLSPTYTGRRHTIVDSKLEIELQGKKYRDSAFIKSGIDWM
VcrA          121  QHKGVYEG--PPD-------APFTSWGNR----LQVRMSGEEQKKRI---------LAAK
BvcA          121  NHQGAYWGPVPEN-------GDDKRYPDP----ADQCLTLPEKRDLY---------LAWA
TcbA          121  SKITGKKDT-----------GKDLPTLNAERLGIKGRPATNTETSVL-FYSQHLGAMLTQ
TmrA          121  RWNG------------------------------------------------------WQ
CfrA          121  RWNG------------------------------------------------------WQ
AcdA          121  RWNG------------------------------------------------------WQ
```

```
NpRdhA       181 ---------------G-FPARGHFSGDAQVDLARLAVRAGLARVVDGVLVAPFLRRGFRL
PceA (Sfs)   181 -------------ENG-EN--------------------GPKMKAKNVGEARAGRALEA-
PceA (Dcc)   181 DKKAEDEILYAKARED-FP--------------------GWEPGLDGFGD----IRTTAL
PceA (Dhb)   181 -------------RHN-ET--------------------GWT-GLDEALN----AGAWA-
TceA         181 -------------KENIDP--------------------DYDPGELGYGD----RREDA-
VcrA         181 -------------KER-FP--------------------GWDGGLHGRGD----QRADA-
BvcA         181 -------------KQQ-FP--------------------DWEPGINGHGP----TRDEA-
TcbA         181 -------------RHN-ET--------------------GWT-GLDEALN----AGAWA-
TmrA         181 -------------TDG-SP--------------------GLT-VLDGAAA----RASFA-
CfrA         181 -------------TDG-SP--------------------GLT-VLDGAAA----RASFA-
AcdA         181 -------------TDG-SP--------------------GLT-VLDGAAA----RASFA-

NpRdhA       241 GVVTTGYALAADRPLAPEGD--------LGETAPEVMLGIDGTRP----GWEDAE-EEKR
PceA (Sfs)   241 ----AGWTLDINYGNIYPNR--FFMLW-SGETMTNTQ------------LWAPVG-----
PceA (Dcc)   241 THASEMFSFGNFPTRMNLGG--NMVDLVAAVRAAGGYLGSTD-------SYAGPK-MVHT
PceA (Dhb)   241 ----VEFDYSGFNAAGGGPG--SVIPL-YPINPMTNEIANEPVMVPGLYNWDNID---VE
TceA         241 ----LIYAATNGSHNCWENPLYGRYEGSRPYLSMRTMNGINGLH-----EFGHAD-----
VcrA         241 ----LFYAVTQPFPGSGEEG-HGLFQP-YPDQPGKFYAR-----------WGLYGPPHDS
BvcA         241 ----LWFASSTGG--------IGRYRI-PGTQQMMSTMRLDGSTG----GWGYFN-QPPA
TcbA         241 ----VEVDYSGFNAAGGGPG--SVITP-YPINPMTNEIANEPVMVPGLYNWDNID---VE
TmrA         241 ----VDYYFNGENSACRANK--GFFEW-HPKVPELNF------------KWGDPE----R
CfrA         241 ----VDYYFNGENSACRANK--GFFEW-HPKVAELNF------------KWGDPE----R
AcdA         241 ----VDYYLNGENSACRANK--GFFEW-HPKVPELNF------------KWGDPE----R

NpRdhA       301 PLHMGRYPMETIRRVDEPTTLVVRQEIQRVAKRGDFFKRAEAGDLGEKAKQEKKRFPMKH
PceA (Sfs)   301 ---LDRRPPDT--TDPVELTNYVKFAAR-------MAGADLVGVARLNR----N------
PceA (Dcc)   301 PEEMGGTKYQ---GTPEDNLRTLKAGIR-------YFGGEDVGALELDD-KLKK------
PceA (Dhb)   301 SVRQQGQQWKF--ESKEEASKILKKATR-------LLGADLVGIAPYDE----R------
TceA         301 -IKTTNYPKWE--GTPEENLLIMRTAAR-------YFGASSVGAIKITD-NVKK------
VcrA         301 APPDGSVPKWE--GTPEDNFLMLRAAAK-------YFGAGGVGALNLADPKCKK------
BvcA         301 AVWGGKYPRWE--GTPEENTLMMRTVCQ-------FFGYSSIGVMPITS-NTKK------
TcbA         301 SVRQQGQQWKF--KSKEEASKMVKKAAC-------FLGADLAGIAPYDE----R------
TmrA         301 NIHSPGV------KSAEEGTMAVKRMAR-------FFGAAKAGIAPFDK----R------
CfrA         301 NIHSPGV------KSAEEGTMAVKKIAR-------FFGAAKAGIAPFDK----R------
AcdA         301 NIHSPGV------KSAEEGTMAVKKIAR-------FFGAAKAGIAPFDK----R------

NpRdhA       361 PLALGMQPLIQNMVPLQGTREKL---AP--TGKGGDLSDPGRNAEAIKALGYYL------
PceA (Sfs)   361 ------------WVYSEAVTI------PADVPYEQSLHKEIEKPIVFKDVPLPI------
PceA (Dcc)   361 ------------LIFTV-------------DQYGKALEFGDVEECIETPKQVTI------
PceA (Dhb)   361 ------------WTYSTWGRKIL---KPCKMPNGRT-KY----------LPWDLPKMLSG
TceA         361 ------------IFYAKVQPFCL-GPWYTITNMAEYIEYPV--PVDNYAIPIVFEDIPAD
VcrA         361 ------------LIYKKAQPMTLGKGTYSEIGGPGMIDAKIYPKVPDHAVPINFKEA--D
BvcA         361 ------------LFFEKQIPFQFMAGDPGVFGGTGNVQFDV--PLPKTPVPIVWEEV--D
TcbA         361 ------------WTYSTWGRKIL---KPCKMPNGRT-KL----------MPWDLPKVLSG
TmrA         361 ------------WVFTETAAFVK-------TPEGEDLKF----------IPPDF------
CfrA         361 ------------WVFTETYAFVK-------TPEGESLKF----------IPPDF------
AcdA         361 ------------WVFTETAAFVK-------TPEGESLKF----------IPPDF------
```

```
NpRdhA       421 -GADFVGICRAEP-WMYYASDEVEGKPIEAYHDYAVVMLIDQGYETMEGASGDDWISASQ
PceA (Sfs)   421 -----------ETDDELIIPNTCE---------NVIVAGIAMNREMMQTAP--NSMACAT
PceA (Dcc)   421 -------------------PNKCK---------YIFLWTMRQPYEWTRRQS--GRFEGAA
PceA (Dhb)   421 GGVEVFGHAKFEPDWEKYAGFKPK---------SVIVFVLEEDYEAIRTSP--SVISSAT
TceA         421 QGHYS--YKRFGGDDKIAVPNALD---------NIFTYTIMLPEKRFKYAH--SIPMDPC
VcrA         421 YSYYN------DAEW--VIPTKCE---------SIFTFTLPQPQELNKRTG--GIA-GAG
BvcA         421 KGYYN--------DQKIVIPNKAN---------WVLTMTMPLPEDRFKRSL--GWSLDAS
TcbA         421 GGVEVFGHAKFEPDWEKYAGFKPK---------SVIVFVLEEDYEAIRTSP--SVISSAT
TmrA         421 -------------------GFEPK---------HVISMIIPQSLEGVKCAP--SFLGSAE
CfrA         421 -------------------GFEPK---------HVISMIIPQSPEGVKCDP--SFLGSTE
AcdA         421 -------------------GFEPK---------HVISMIIPQSLEGIKCAP--SFLGSAE

NpRdhA       481 SMRAYMRGAEIAGVMAAHCRRMGYSARSHSNA-HSEVIHNPAILMAGLGEVSRIGDTLLN
PceA (Sfs)   481 TAFCYSRMCMFDMWLCQFIRYMGYYAIPSCNG-VGQSV--AFAVEAGLGQASRMG-ACIT
PceA (Dcc)   481 TETSYERAYNTKAHFQDFVRGLGYQMISAGNNSLSPAG--AWAVLGGLGELSRAS-YVNH
PceA (Dhb)   481 VGKSYSNMAEVAYKIAVFLRKLGYYAAPCGND-TGISV--PMAVQAGLGEAGRNG-LLIT
TceA         481 SCIAYPLFTEVEARIQQFIAGLGYNSMGGGVEAWGPGS--AFGNLSGLGEQSRVS-SIIE
VcrA         481 SYTVYKDFARVGTLVQMFIKYLGYHALYWPIG-WGPGG--CFTTFDGQGEQGRTG-AAIH
BvcA         481 SMIAYPQMAFNGGRVQTFLKALGYQGLGGDVAMWGPGG--AFGVMSGLSEQGRAA-NEIS
TcbA         481 LGKSYSNMAEVAYKIAVFLRKLGYYAAPCGND-TGINV--PMAVQAGLGEAGRNG-LLIT
TmrA         481 YGLSFAQIGYAAFGLSMFIKDLGYHAVPIGSD-SALSI--PIAIQAGLGEYSRSG-QMIT
CfrA         481 YGLSCAQIGYAAFGLSMFIKDLGYHAVPIGSD-SALAI--PIAIQAGLGEYSRSG-LMIT
AcdA         481 YGLSYAQIGYAAFGLSMFIKDLGYHAVPIGAD-SALAV--PIAIQAGLGEYSRSG-LMIT

NpRdhA       541 PFIGPRSKSIV-FTTDLPMSVDRPIDFGLQDFCNQCRKCARECPCNAISFG-DKVMFN--
PceA (Sfs)   541 PEFGPNVRLTK-VFTNMPLVPDKPIDFGVTEFCETCKKCARECPSKAITEG--PRTFEGR
PceA (Dcc)   541 PLYGITVRVTWGFLTDMPLPPSRPIDFGARKFCETCGICAENCPFGAINPG-EPTWKDDN
PceA (Dhb)   541 QKFGPRHRIAK-VYTDLELAPDKPRKFGVREFCRLCKKCADACPAQAISHEKDPKVLQPE
TceA         541 PRYGSNTKGSLRMLTDLPLAPTKPIDAGIREFCKTCGICAEHCPTQAISHE-GPRYDSPH
VcrA         541 WKFGSSQRGSERVITDLPIAPTPPIDAGMFEFCKTCYICRDVCVSGGVHQEDEPTWDSGN
BvcA         541 PKYGSATKGSNRLVCDLPMVPTKPIDAGIHKFCETCGICTTVCPSNAIQVG-PPQWSNNR
TcbA         541 QKFGPRHRIAK-VYTDLELAPDKPRKFGVREFCRLCKKCADACPAQAISHEKDPKVLQPE
TmrA         541 PEFGPNVRLCE-VFTDMPLNHDKPISFGVTEFCKTCKKCAEACPPQAISYE-DPTIDGPR
CfrA         541 PEFGSNVRLCE-VFTDMPLNHDKPISFGVTEFCKTCKKCAEACAPQAISYE-DPTIDGPR
AcdA         541 PEFGSNVRLCE-VFTDMPLNHDKPISFGVTEFCKTCKKCAEACPPQAISYE-DPTIDGPR

NpRdhA       601 --------GYEIWKADVEKCTKYRVTQMKGSACGRCMKMCPW----NREDTVEGRRLAEL
PceA (Sfs)   601 S--IHNQSGKLQWQNDYNKCLGYWPES--GGYCGVCVAVCPF----TKGNIWIHDGVEWL
PceA (Dcc)   601 ---AFGNPGFLGWRCDYTKCPH----------CPICQGTCPF---NSHPGSFIHDVVKGT
PceA (Dhb)   601 DCEVAENPYTEKWHLDSNRCGSFWAYN--GSPCANCVAVCSW----NKVETWNHDVARIA
TceA         601 ---WDCVSGYEGWHLDYHKCIN----------CTICEAVCPF--FTMSNNSWVHNLVKST
VcrA         601 ---WWNVQGYLGYRTDWSGCHNQ---------CGMCQSSCPFTYLGLENASLVHKIVKGV
BvcA         601 ---WDNTPGYLGYRLNWGRCVL----------CTNCETYCPF--FNMTNGSLIHNVVRST
TcbA         601 DCEVAENPYTEKWHLDSNRCGSFWAYN--GSPCSNCVAVCSW----NKVETWNHDVARIA
TmrA         601 G--QMHNSGIKRWYVDPVKCFEFWSRDNVRNCCGACIAACPF----TKPEAWHHTLIRSL
CfrA         601 G--QMQNSGIKRWYVDPVKCLEFMSRDNVGNCCGACIAACPF----TKPEAWHHTLIRSL
AcdA         601 G--QMHNSGIKRWYVDPVKCFEFWSRDNVRNCCGACIAACPF----TKPEAWHHTLIRSL
```

```
NpRdhA       661 SIKVPEARAAIIAMDDALQNGKRNLIKRWWFDLEVIDGVAGAPRMGTNERDLSPDRGDKI
PceA (Sfs)   661 IDNTRFLDPLMLGMDDALGYGAKRNITEVWDGKINTYGLD--------------------
PceA (Dcc)   661 VSTTPIFNSFFKNMEKTFKYGRKNP-----------------------------------
PceA (Dhb)   661 T-QIPLLQDAARKFDEWFGYNGPVNPDERLESGYVQNMV---------------------
TceA         661 VATTPVFNGFFKNMEGAFGYGPRYSPSR--------------------------------
VcrA         661 VANTTVFNSFFTNMEKALGYGDLTMEN---------------------------------
BvcA         661 VAATPVFNSFFRQMEHTFGYGMKDDL----------------------------------
TcbA         661 T-RIPLLQDAARKFDEWFGYNGPVNPEERLESGYVQNMV---------------------
TmrA         661 V-GAPVITPFMKDMDDIFGYGKPNEKAK--------------------------------
CfrA         661 V-GAPVITPFMKDMDDIFGYGKLNDEKAI-------------------------------
AcdA         661 V-GAPVITPFMKDVDDIFGYGKPNDEKAI-------------------------------

NpRdhA       721 GANQKLAMYPPRLQPPPGTTLDAVLPVDRSGGLAEYAAAETPAA-ARARLKSSAG---
PceA (Sfs)   721 ---------------------------------ADH------FR-DTVSFRKDRVKKS
PceA (Dcc)   721 ---------------------------------ATW------WDEVDDYPYGVDTSY-
PceA (Dhb)   721 ---------------------------------KDF------WN-NPESIKQ------
TceA         721 ---------------------------------DEW------WA-SENPIRGASVDIF
VcrA         721 ---------------------------------SNW------WK-EEGPIYGFDPGT-
BvcA         721 ---------------------------------NDW------WNQSHKPW--------
TcbA         721 ---------------------------------TDF------WN-NPESIKQ------
TmrA         721 ---------------------------------ADW------WK--------------
CfrA         721 ---------------------------------ADW------WK--------------
AcdA         721 ---------------------------------ADW------WK--------------
```
